# The superior colliculus response space has globally high– and locally low-dimensionality

**DOI:** 10.1101/2023.11.06.565916

**Authors:** Ole S. Schwartz, Keisuke Yonehara

## Abstract

An understanding of cell types is essential for understanding neural circuits, but only when the response of each type is clearly defined and predictable, as has been observed in the retina^1^. Recent work has shown that neural responses in the visual cortex are of high dimensionality, questioning the validity of defining cell types in the deeper visual system^2–4^. Here we investigate the dimensionality of neural responses in the midbrain using two-photon calcium imaging in superficial layers of the mouse superior colliculus (SC). Responses of individual neurons to closely related stimuli, such as ON and OFF light signals, were mutually dependent such that the response to one stimulus could be predicted from the response to the other. In contrast, individual neurons responded to brightness and motion in a statistically independent manner, maximizing functional diversity but preventing traditional cell type classification. To capture the globally high, locally low dimensionality of neural responses, we propose a multidimensional response model, in which classification of cellular responses is meaningful only in local low-dimensional structures. Our study provides a framework to investigate the processing of visual information by the SC, which likely requires a high-dimensional response space structure^5,6^ to perform higher-order cognitive tasks^7–12^.

## Introduction

Classification of neurons according to genetic and functional types has been a powerful way to understand the functional organization of the nervous system^1^, enabling targeted recordings and genetic therapies. The goal of cell type classification has been to uncover a single set of types, such that each cell belongs to only one type^1^, and has relied on the assumption that a meta-structure covering all response features exists. This approach works well in a low dimensional response space, where features are correlated such that classification based on feature x will be similar to classification based on feature y. However, in a high dimensional response space, features are uncorrelated, preventing compression into a single classification.

Classification of neurons in the retina has been relatively successful^13–17^. Retinal ganglion cells (RGCs) have functional, morphological, and molecular features that are well matched^13,18,19^, including OnOff direction-selective cells tuned to fast visual stimuli and On direction-selective cells tuned to slow stimuli. These cell-type-specific responses are further defined by the depth of dendritic stratification and synaptic specificity within the inner plexiform layers, according to the complement of genes expressed. Together, these findings suggest a low-dimensional organization of retinal response space. In contrast, cell-type classification has been less effective in downstream visual areas^20–22^, including the visual cortex, where response space has a relatively high dimensionality^2,6^.

The SC is an ancient visual structure in the midbrain that can be divided into a superficial region (sSC) that relays visual information to the cortex and a deep multimodal part (dSC) that transforms sensory information into motor commands^7,23–28^. In mice, it receives retinotopically organized input from ∼90% of RGCs^23,29^, of which there are ∼40 different types^13^. However, despite several attempts at creating a functional classification system for the SC, only 4-5 distinct morpho-physiological types have been identified^20,21,30,31,32^. Moreover, the question of response space dimensionality in the SC has not been addressed.

We investigated the dimensionality of response space in the SC and examined whether this dimensionality influences the ability to functionally classify neurons. Using two-photon calcium imaging of the SC in awake mice, we recorded the responses of 6,872 neurons to different light stimuli. An overall statistical independence between responses to unrelated stimuli, as well as mutual dependence between responses to related stimuli, suggested that the SC response space is multidimensional. Moreover, this multidimensionality limited the ability to cluster neurons according to cell type. Our results provide a conceptual framework for functional classification in high-dimensional response spaces as well as insight into how retinal information is transformed at the first central visual structure.

## Results

### Unrelated stimuli drive independent responses

To investigate visual response space in the midbrain, we collected functional responses of sSC neurons to four visual stimuli that elicit a range of visual responses (Figure 1a-b, Supplementary Figure 1). The stimuli included: 1) A 10-degree circular chirp-modulated stimulus (chirp); 2) A full-screen sinusoidal grating drifting at 5 degrees per second (d/s) in 12 directions (slow drift); 3) A full-screen sinusoidal grating drifting at 40 d/s in 12 directions (fast drift); and 4) A full-screen sinusoidal grating drifting in all combinations of 6 temporal and 8 spatial frequencies in 4 directions (SpaTemp). We included the chirp stimulus to identify cell types that can detect increases (ON) and decreases (OFF) in luminance, the drift stimuli to identify cell types that have orientation and/or direction selectivity^13,21,32^, and the SpaTemp stimulus to determine the speed preference of cells.

**Figure 1.**
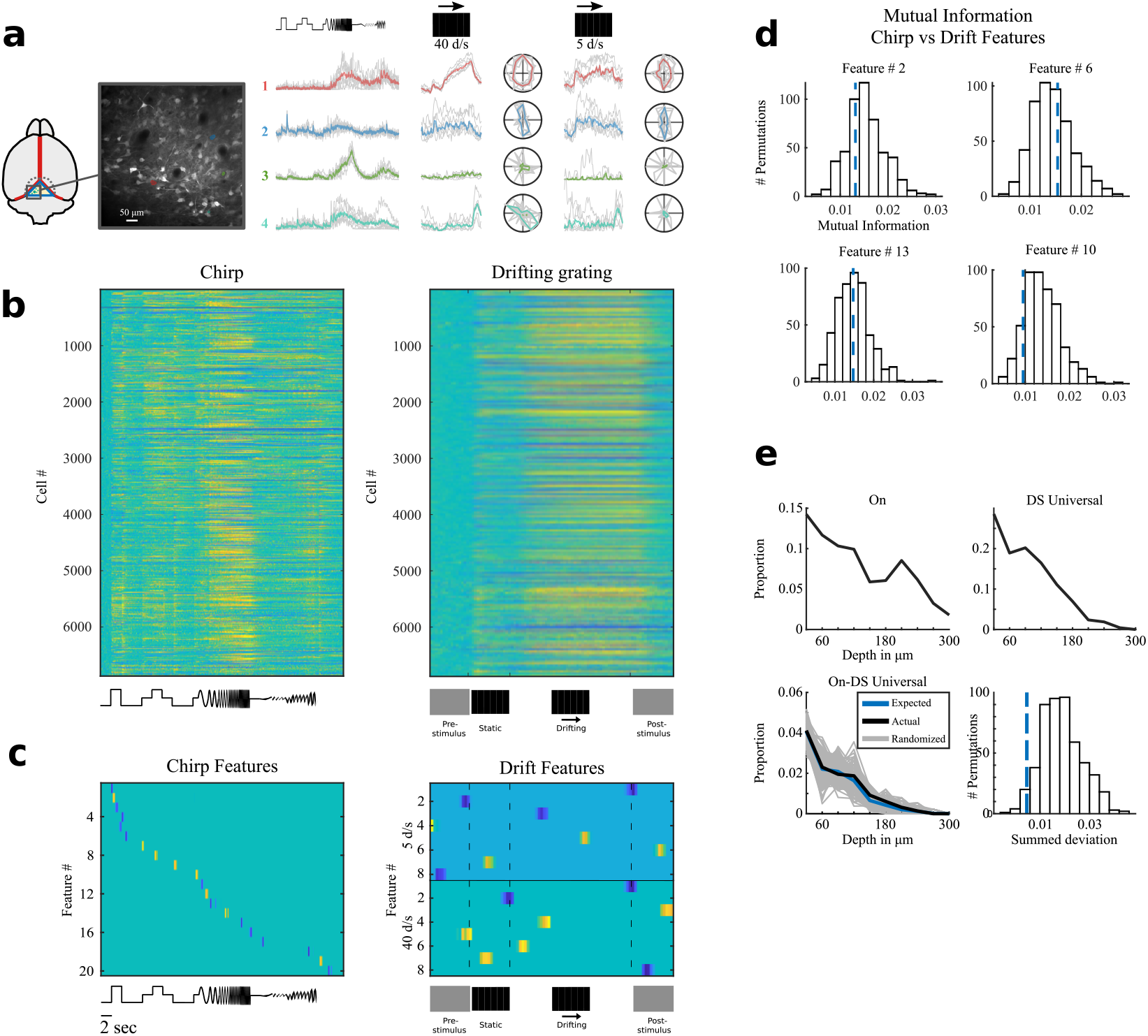
Global Independencies. **a**) Left: Illustration of mouse brain with silicone plug (blue), displacing sinuses (red) to create accessible visual field of superior colliculus cells. Middle: zoom-in of a field of view in two-photon microscope. Right: traces of four cells (outlined on middle image) responding to changes in luminance of the chirp stimulus and moving gratings at two different speeds. **b)** Response matrices for the chirp (left) and slow grating (right) stimuli. **c)** Matrices showing the weight of each chirp (left) and drift (right) feature. Blue indicates negative weight and yellow indicates positive weight. **d)** Mutual information for four example chirp features vs the drift feature with which they had the highest MI (dashed blue lines) plotted on top of histograms of 500 calculations of MI after the order of the drift feature had been randomized relative to the order of the chirp feature. **e)** Example of a predictable relationship between response types. Top left: Proportion of On cells across depth. Top right: Proportion of DS Universal cells across depth. Bottom left: Actual On-DS Universal proportions across depth (black) vs expected proportion (blue) and random permutations (gray). Bottom right: Histogram of the summed deviation from the expected proportion of each random permutation with the actual summed deviation shown as a dashed blue line.

We extracted 20 features from the responses to the chirp stimulus using sparse principal component analysis (sPCA) and 16 features from the responses to the two drift stimuli using singular value decomposition followed by sPCA (Figure 1c, Supplementary Figure 2-3). Mutual information (MI) ^33^ was then employed to evaluate the degree of dependency between the drift and chirp features (see Methods). If features x and y have high MI, feature x can provide information about the nature of feature y in a given cell. However, if there is no or chance MI between two features, any overlap between two features in a given cell is coincidental, and the fraction of cells with a specific combination of these features is predictable based on the distribution of two features across the cell population.

To test the degree of dependency between responses to drift and chirp, we calculated MI between all combinations of the 20 chirp features and the 16 drift features and found chance MI in all cases (320 permutation tests, Wilson’s harmonic mean p = 0.11, lowest p-value after Bonferroni-Holm correction = 0.64, Supplementary Figure 4). MI between chirp and SpaTemp features was also determined to be at chance level in all cases (100 permutation tests, Wilson’s harmonic mean p = 0.27, all p-values > 1 after Bonferroni-Holm correction). These results suggest that the shape of a cell’s response to the chirp stimulus is not predictive of its response to either of the drift or the SpaTemp stimuli.

We also manually extracted ON/OFF response amplitudes to classify cells as On, Off, or OnOff, and calculated orientation and direction selectivity indexes (OSi and DSi, respectively) to classify cells as orientation-selective (OS) or direction-selective (DS) (Supplementary Figure 5A-B). We further quantified the response to luminance changes by measuring how fast the peak amplitude was reached (time-to-peak) and how sustained the response was (min-to-peak ratio). Furthermore, because we used drift stimuli with two different speeds, we could subdivide DS cells into DS Fast (direction-selective only to the fast grating), DS Slow (direction-selective only to the slow grating), and DS Universal (direction-selective at both speeds). MI between ON/OFF response amplitudes and manually defined OS and DS features (DSi, OSi, preferred orientation, and preferred direction) was at chance-level (16 Wilcoxon’s rank sum tests, Wilson’s harmonic mean p = 0.32, all p-values > 1 after Bonferroni-Holm correction; Supplementary Figure 6). Thus, responses to brightness cannot predict responses to motion, revealing the independence of responses to unrelated stimuli in the SC.

If two independent groups exist in the same space, they will overlap predictably (Supplementary Figure 7). To test if collicular On/Off and OS/DS response types are independent, we investigated whether the distribution of the combined groups (e.g. On cells that are also DS Universal) could be predicted from the depth distribution of their separate groups (e.g. On cells and DS Universal cells; Figure 1e). For all combinations of On/Off and OS/DS subtypes, we found the overlap to be predictable (Supplementary Figure 8, 12 permutation tests, Wilson’s harmonic mean p = 0.08, lowest p-value = 0.10 after Bonferroni-Holm correction).

Having found that all overlaps between On/Off and OS/DS subtypes are predictable, we reasoned that the response properties of the combined groups should not differ from the response properties of the remaining cells in the two parent groups. We therefore calculated the distribution of min-to-peak ratio, time-to-peak, and preferred orientation and direction for all 12 subtype combinations and confirmed that the distribution of response properties of subtype combinations (e.g., On-DS Universal cells) did not differ from that of the remaining cells in their parent group (e.g., On-nonDS Universal cells) in all cases except the On-DS Slow combination, whose response properties deviated less than chance from the expected distribution (32 Wilcoxon’s rank sum tests for distribution of ON/OFF response properties, Wilson’s harmonic mean p = 0.21, all p-values > 1 after Bonferroni-Holm correction; 12 t-tests for distribution of OS/DS response properties, p = 0.01 for On-DS Slow after Bonferroni-Holm correction, Wilson’s harmonic mean p = 0.27 for all other combinations, lowest p-value = 0.74 after Bonferroni-Holm correction; Supplementary Figure 8).

We confirmed this observation by subdividing the response types into 12 On/Off subtypes (each On, Off and OnOff cell classified by sustained, transient, fast or slow responses) and 14 OS/DS subtypes (each DS Slow, DS Fast, DS Universal, and OS cell classified by the cardinal directions). After performing the same tests as described above, we found that the proportion of cells and distribution of response properties could be predicted from those of the parent groups for all 168 combinations (168 t-tests, 396 Wilcoxon’s rank sum tests, Wilson’s harmonic mean p = 0.2, all p-values > 1 after Bonferroni-Holm correction).

### Chirp and drift clusters overlap at random

A commonly used strategy to identify clusters of cellular responses is to use a range of stimuli that cover the largest possible stimulus space^13,21^. However, such a strategy may not be feasible when there are statistical independencies within the response space, as independent features reduce the ability of the clustering algorithm to separate response types and increase the risk of mistaking coincidental overlap for cell types (Supplementary Figure 7). We tested the ability to identify clusters in the SC by performing gaussian mixture model clustering of response features to chirp and drift, both individually and collectively. We identified 50 clusters for drift stimuli, 28 clusters for chirp stimuli, and 31 clusters for both chirp and drift stimuli (Supplementary Figure 9A-C, 10A-C). All clusters were stable (median Jaccard similarity score = 0.55 for chirp, 0.56 for drift, and 0.47 for chirp-drift).

As expected from the statistical independence between On/Off and OS/DS response features (Figure 1), we found only chance overlap between chirp-derived clusters and drift-derived clusters (permutation test, p = 0.76; Figure 2a). Chirp and drift clustering is successful at separating On/Off and OS/DS response types, respectively, but combined chirp-drift clustering performs worse at separating these response types (Figure 2b-c). Furthermore, drift clustering cannot separate On/Off response types better than chance (permutation tests, p = 0.95 for On, 0.33 for Off, and 0.98 for OnOff), and chirp clustering cannot separate OS/DS response types better than chance (permutation tests versus random data, p = 0.67 for DS Slow, 0.62 for DS Fast, 0.91 for DS Universal, and 0.21 for OS; Supplementary Figure 11).

**Figure 2.**
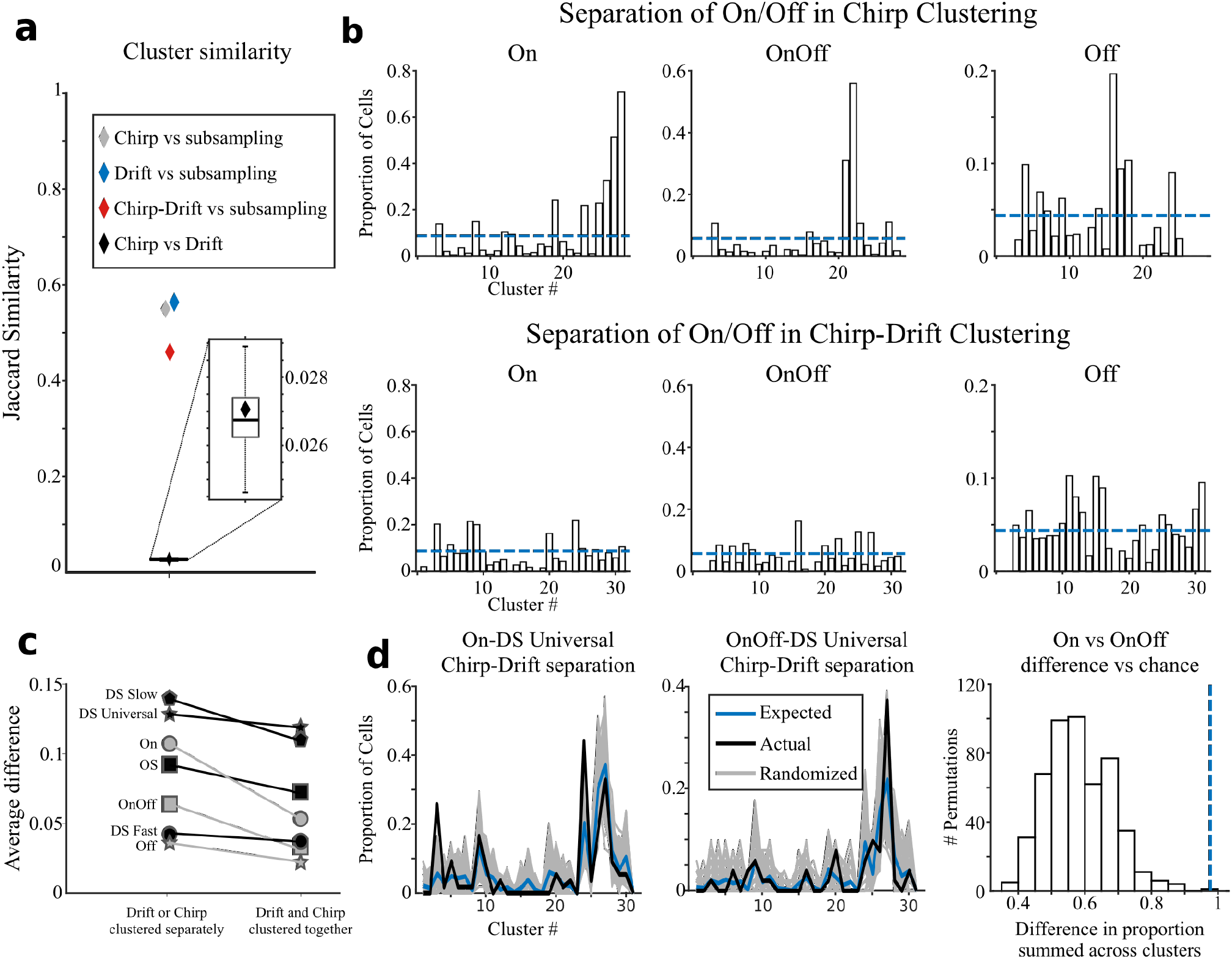
Clustering Independencies. **a**) Median Jaccard Similarity score for chirp compared to 90% chirp subsamples (grey diamond), for drift compared to 90 % drift subsamples (blue diamond), and for chirp clustering compared to drift clustering (black diamond). Boxplot shows median Jaccard Similarity score for chirp clusters compared to 500 random permutations of the drift cluster tags. **b)** Proportion across clusters of On (left), OnOff (middle), and Off (right) for the chirp clustering (top) and chirp-drift clustering (bottom). Dashed blue line indicates expected value if distribution is random. **c)** A clustering’s ability to separate On/Off and OS/DS response types can be quantified as the absolute difference between the actual and the expected proportion of the given response type averaged across clusters. For all measured response types, the ability to separate types is lower in the combined clustering (right) than in the separate clusterings (left). **d)** Left: Actual distribution of On-DS Universal cells across clusters (black) is different from expected distribution if sampling is random (blue, p = 1.8e-6). 500 random permutations are shown in gray. Middle: Same as left for the OnOff-DS Universal population (p = 0.0051). Right: Sum of the absolute difference between On-DS Universal cluster proportions and OnOff-DS Universal cluster proportions (dashed blue line, p = 6.6e-5) plotted on top of a histogram of 500 sums of the absolute difference between a random sample of On cells of equal size and depth distribution as On-DS Universal cells (gray lines in left figure) and a random sample of On cells of equal size and depth distribution as OnOff-DS Universal cells (gray lines in middle figure).

Although On/Off and OS/DS response types are statistically independent and the response properties of the combined groups do not differ from those of the parent groups, combined chirp-drift clustering separate OS/DS responses into On, Off, and On-Off subtypes (Figure 2d). Taken together, these results show an absence of overlap between clusters based on chirp and drift features and that combining statistically independent features into a single clustering does a worse job of separating response types than clustering these features independently. Moreover, clustering based on the combination of statistically independent features creates meaningless subdivisions of response types based on coincidental overlap between responses. Indeed, including drift features in chirp clustering, and vice versa, disrupts the clustering more than adding a similar amount of noise suggesting that the structures uncovered by drift and chirp are orthogonal (p = 0.02 for adding drift to chirp vs noise to chirp, p = 0.02 for adding chirp to drift vs adding noise to drift; Supplementary Figure 12).

### Related stimuli can result in local dependencies

If On/Off and OS/DS responses are independent in the SC, but dependent in the retina, it is possible that the response space structure of the SC is closer to that of the cortex than the retina. Cortical response space is thought to be high-dimensional at the global scale but differentiable for related stimuli at a local scale^2^. We therefore anticipated local dependencies in the SC despite overall statistical independence, which we tested by examining dependencies within responses to chirp and drift. MI between ON and OFF responses to the chirp stimulus was higher than chance (permutation test, p < 1 x 10^−4^). Consistent with the high MI we also found that the proportion of OnOff responses by depth could not be explained as coincidental overlap between significant ON and OFF responses (t-test versus random permutations, p = 3.5 x 10^−124^; Figure 3a).

**Figure 3.**
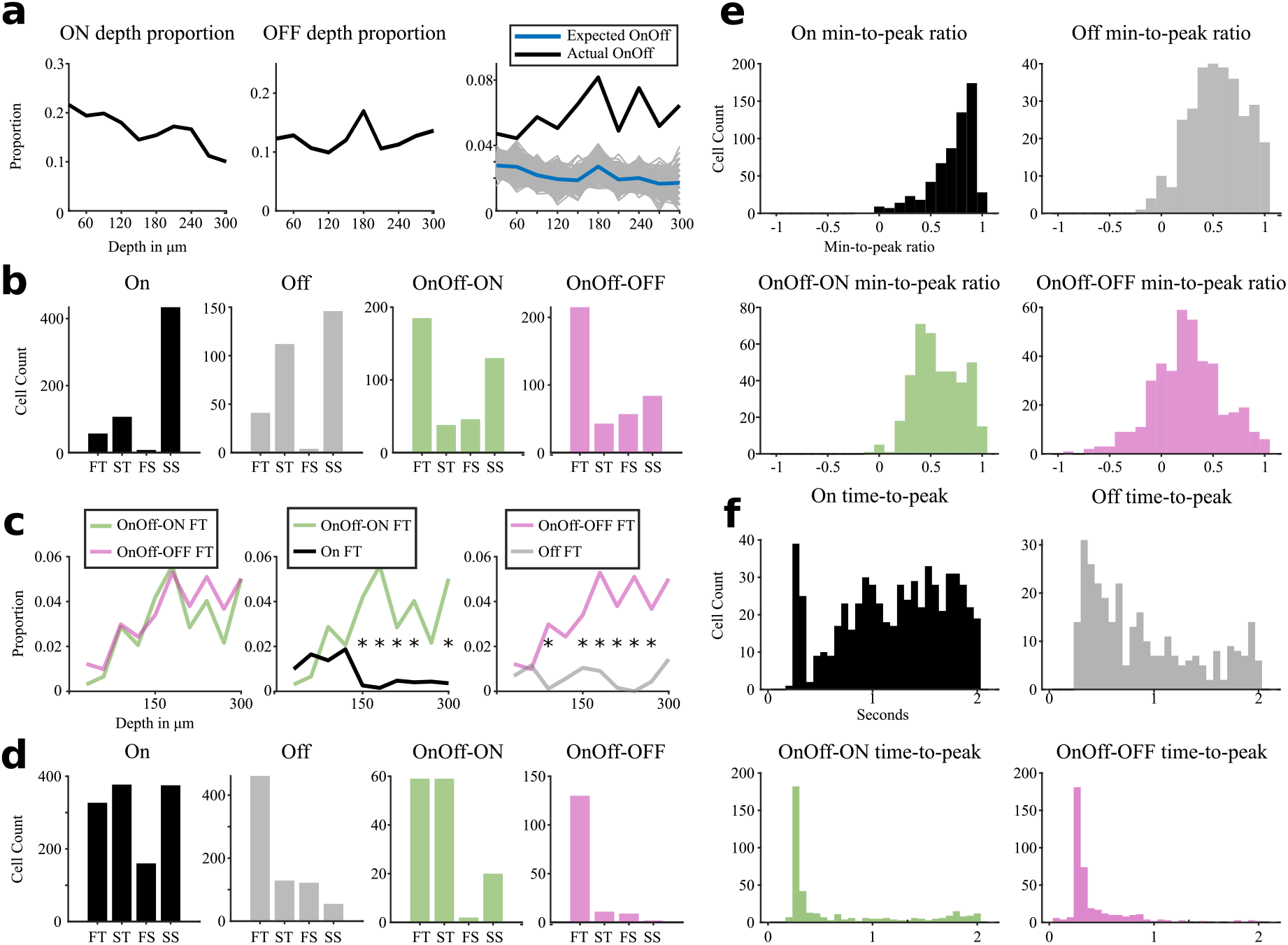
Local dependencies in ON/OFF Response Space. **a**) Proportion of above threshold ON (left) and OFF responses (middle) across depth. Right: Actual proportion of OnOff cells across depth (black) compared to expected proportions if ON-OFF overlap was coincidental (blue) with 500 random permutations of linear combinations of On and Off shown in gray. **b)** On/Off subtype composition in the SC: Composition of Fast-Transient (FT), Slow-Transient (ST), Fast-Sustained (FS), and Slow-Sustained (SS) subtypes across On, Off, and OnOff cells. Because subtypes can be determined in relation to both the ON and OFF phase, the subtype composition for OnOff cells is shown for each phase separately. **c)** Left: Proportions across depth of OnOff-ON FT cells (green) and OnOff-OFF FT cells (magenta). Middle: Proportions across depth of OnOff-ON FT cells (green) and On FT cells (black). Right: Proportions across depth of OnOff-OFF FT cells (magenta) and Off FT cells (grey). **d)** On/Off subtype composition in the Retina: Shown as in b). Data was extracted from http://retinal-functomics.net/ ^34^. **e)** Distribution of min-to-peak ratio for On (top left), Off (top right), OnOff-ON (bottom left) and OnOff-OFF (bottom right). **f)** Distribution of time-to-peak for On (top left), Off (top right), OnOff-ON (bottom left) and OnOff-OFF (bottom right).

Because min-to-peak ratio and time-to-peak can be extracted from both ON and OFF responses, OnOff cells could be labeled as Transient or Sustained and Fast or Slow in relation to both their ON response (OnOff-ON) and OFF response (OnOff-OFF). We observed that the similarity between the two OnOff response types is higher than the similarity between OnOff-OFF and Off response types and between OnOff-ON and On response types, in relation to both subtype composition (Figure 3b) and anatomical depth (Figure 3c). We also found that min-to-peak ratio (Figure 3e) and time-to-peak (Figure 3f) for OnOff cells were different to those for On or Off cells (Wilcoxon’s rank sum tests, time-to-peak p = 5 x 10^−44^, min-to-peak ratio p = 2 x 10^−22^ for OnOff-ON vs On; time-to-peak p = 1 x 10^−49^, min-to-peak ratio p = 6 x 10^−7^ for OnOff-OFF vs Off). These data are similar to data collected from OnOff RGCs (Figure 3d)^13,34^. In both the retina and SC, OnOff responses are faster and more transient than On and Off responses, and collicular OnOff responses are more similar to retinal OnOff responses than collicular On or Off responses (Figure 3b and d). Our data provide further support for the notion that OnOff is a distinct response type in both the SC and retina.

As expected for locally low dimensionality, we observed high MI between response features extracted from fast-and slow-moving gratings (8 t-tests, 1 x 10^−188^ < p < 1 x 10^−82^ for slow vs best fitting fast features). Indeed, overlap between DS cells responding to the slow grating and DS cells responding to the fast grating was much higher than chance (t-test vs random permutations, p = 5.5 x 10^−136^; Figure 4a). The high MI prompted us to test if there were differences in the preferred direction profile of DS Fast, DS Slow, and DS Universal cells which we found to be the case (Kuiper’s test, Universal vs Fast p < 0.001, Universal vs Slow p < 0.001; Figure 4b). Furthermore, these preferred direction profiles showed that DS cells in the SC segregate into the four cardinal directions. By plotting the change in proportion of preferred direction with depth, we revealed that each cardinal changes independently with depth and that these changes differ between DS subtypes (Figure 4c). Furthermore, the SpaTemp stimulus allowed us to estimate the preferred speed of each cell (see Methods), which differed between cardinals and also changed independently with depth (Figure 4d).

**Figure 4.**
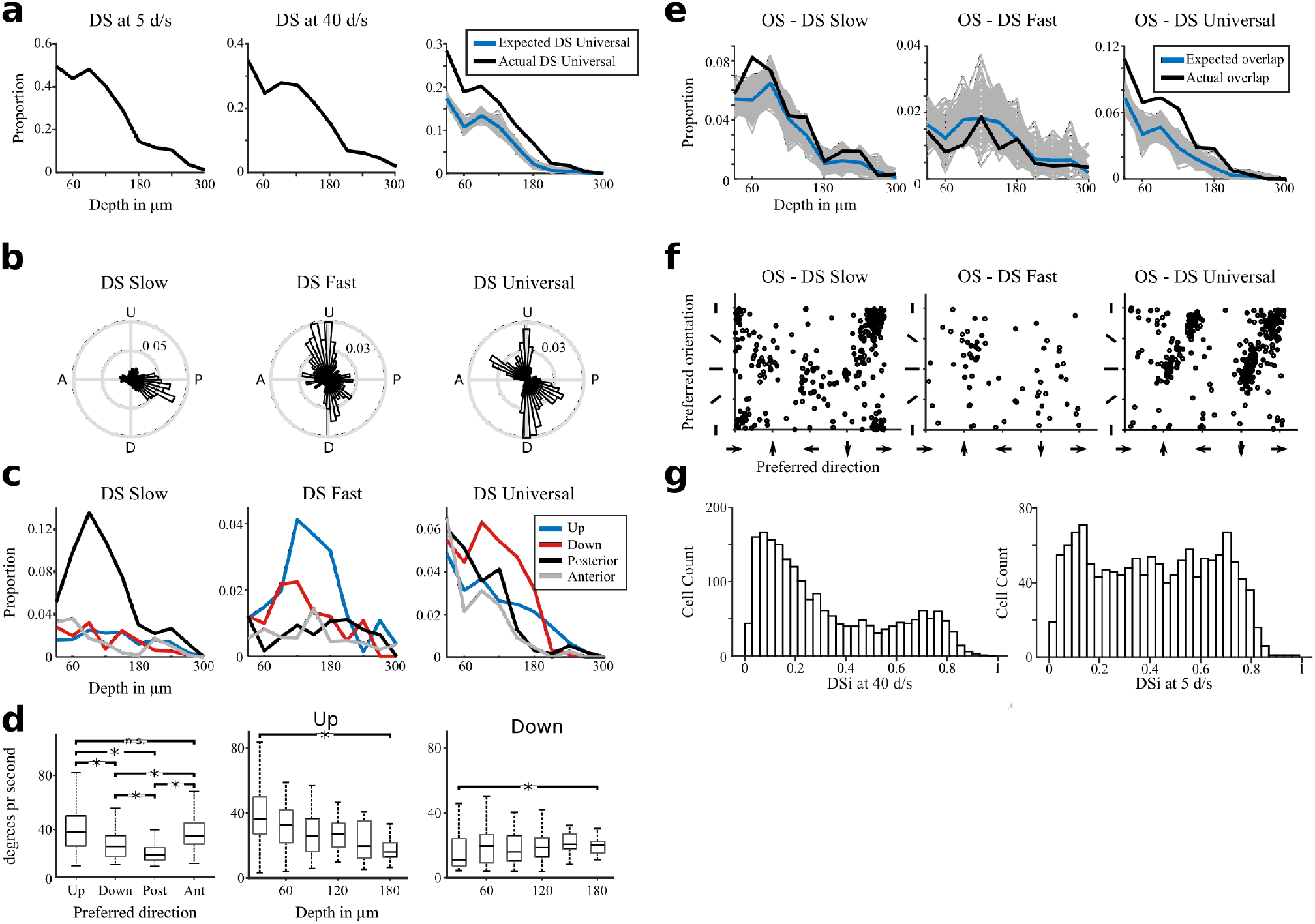
Local Dependencies in OS/DS Response Space. **a**) Proportion of cells with a direction selective response at 5 d/s (left) and 40 d/s (middle) across depth. Right: Actual proportion of DS Universal cells across depth (black) compared to expected proportions if DS Universal responses were a linear combination of DS at 5 d/s and DS at 40 d/s proportions (blue) with 500 random permutations of linear combinations shown in gray. **b**) Polar plots of preferred direction in DS Slow (left), DS Fast (middle) and DS Universal cells (right). **c**) Proportions across depth of the four cardinal directions for DS Slow (left), DS Fast (middle), and DS Universal (right). **d**) Left: Box plot of preferred speed across DS Universal cardinals. Upward selective cells preferred higher speeds than Downward (p = 3e-14) and Posterior (p = 2e-35) selective cells. Downward selective cells preferred higher speeds than Posterior selective cells (p = 4e-21) and lower speeds than Anterior selective cells (p = 4e-13). Posterior selective cells preferred lower speeds than Anterior selective cells (p = 6e-36). All tests were Wilcoxon’s Rank Sum tests. Middle: Box plots of preferred speed across the depth of SGS for Upward selective DS Universal cells. Right: Box plots of preferred speed across the depth of SGS for Downward selective DS Universal cells. The preferred speed for Upward selective DS Universal cells decreases across the SGS (30 µm vs 180 µm: p = 1.1e-4) while the preferred speed for Downward selective cells increases across the SGS (30 µm vs 180 µm: p = 0.03). p-values of less than 0.001 are marked by a *. **e**) Proportions across depth of OS-DS Slow (left), OS-DS Fast (middle), and OS-DS Universal (right) shown in black. Expected proportions if OS-DS overlap was coincidental shown in blue with 500 random permutations of linear combinations of OS-DS proportions shown in gray. Overlap is at chance level for DS Fast (p = 0.97), and higher than chance for DS Slow (p = 4e-5) and DS Universal (p = 7e-46). **f**) Scatter plots of preferred direction (x-axis) and preferred orientation (y-axis) for OS-DS Slow (left), OS-DS Fast (middle), and OS-DS Universal (right). **g**) Left: Histogram of DSi measured at 40 d/s for all cells that are DS at 5 d/s. Right: Histogram of DSi measured at 5 d/s for all cells that are DS at 40 d/s.

Although OS and DS RGCs form separate mosaics in the retina^35–37^, there have been reports of both dependence and independence between preferred orientation and direction in the SC^38–40^. We therefore investigated how orientation preference is related to direction preference in the SC and found a clear difference between how the three DS subtypes overlap with OS cells. Specifically, we found a large overlap between OS and DS Universal or DS Slow but only a chance-level overlap between OS and DS Fast (t-tests against random permutations, OS vs DS Universal p = 7 x 10^−46^, OS vs DS Slow p = 3.7 x 10^−5^, OS vs DS Fast p = 0.97; Figure 4e). The differences in degree of overlap between these subtypes was also reflected in the relationship between preferred direction and orientation of cells that are both OS and DS (Figure 4f). In this case, although DS Universal and DS Slow cells prefer orientations that are orthogonal to their directional preference, we found no evidence for a relationship between preferred orientation and direction of DS Fast cells.

Together these findings suggest that any overlap between OS and DS Fast is coincidental, whereas that between OS and DS Slow or DS Universal is not, revealing the ability to classify OS-DS Slow and OS-DS Universal as distinct types.

Finally, we found that when taking cells that are DS to the Slow drift stimulus (DS Slow and DS Universal cells) and plotting their DSi to the Fast drift stimulus, there were two distinct groups, one with a high DSi to the Fast drift stimulus (DS Universal cells) and one with low DSi (DS Slow cells) to the Fast drift stimulus (Figure 4g). Thus, despite the globally high dimensionality of the SC, we observed local low dimensionality in response to both luminance changes and motion.

### Functional responses are organized by depth

The SC is a layered structure whose superficial region can be divided into the stratum griseum superficiale (SGS) and stratum opticum (SO). We investigated whether the statistically independent responses that we had observed might change through these layers by quantifying functional changes by depth using depth co-clustering correlation matrices for drift and chirp clusters (see Methods). Briefly, we estimated the functional similarity between two depths by calculating how often a cell from one depth occurred in the same functional cluster as cells from another depth. We found that depth co-clustering correlation matrices for chirp and drift have the same structure (Figure 5a-b), despite having independent response types, features, and clusters. There was a clear functional difference between the SGS and SO (180–210 μm) in both chirp and drift matrices. Furthermore, cells within both drift and chirp clusters tended to stem from a narrow layer in depth, indicative of smooth functional changes across depth within the SGS and SO. Finally, we found that cells from the superficial SGS (30 μm) and the superficial SO (210 μm) tended to appear in the same clusters (Figure 5c), revealing high functional similarity between these two regions. This furthermore suggested a ‘double drop’ pattern of functionality with depth, comprising a smooth decrease with depth in the SGS (30-180 μm), a sharp increase between the SGS and SO (180-210 μm), and a second smooth decrease throughout the SO (210-300 μm) (Figure 5d-f).

**Figure 5.**
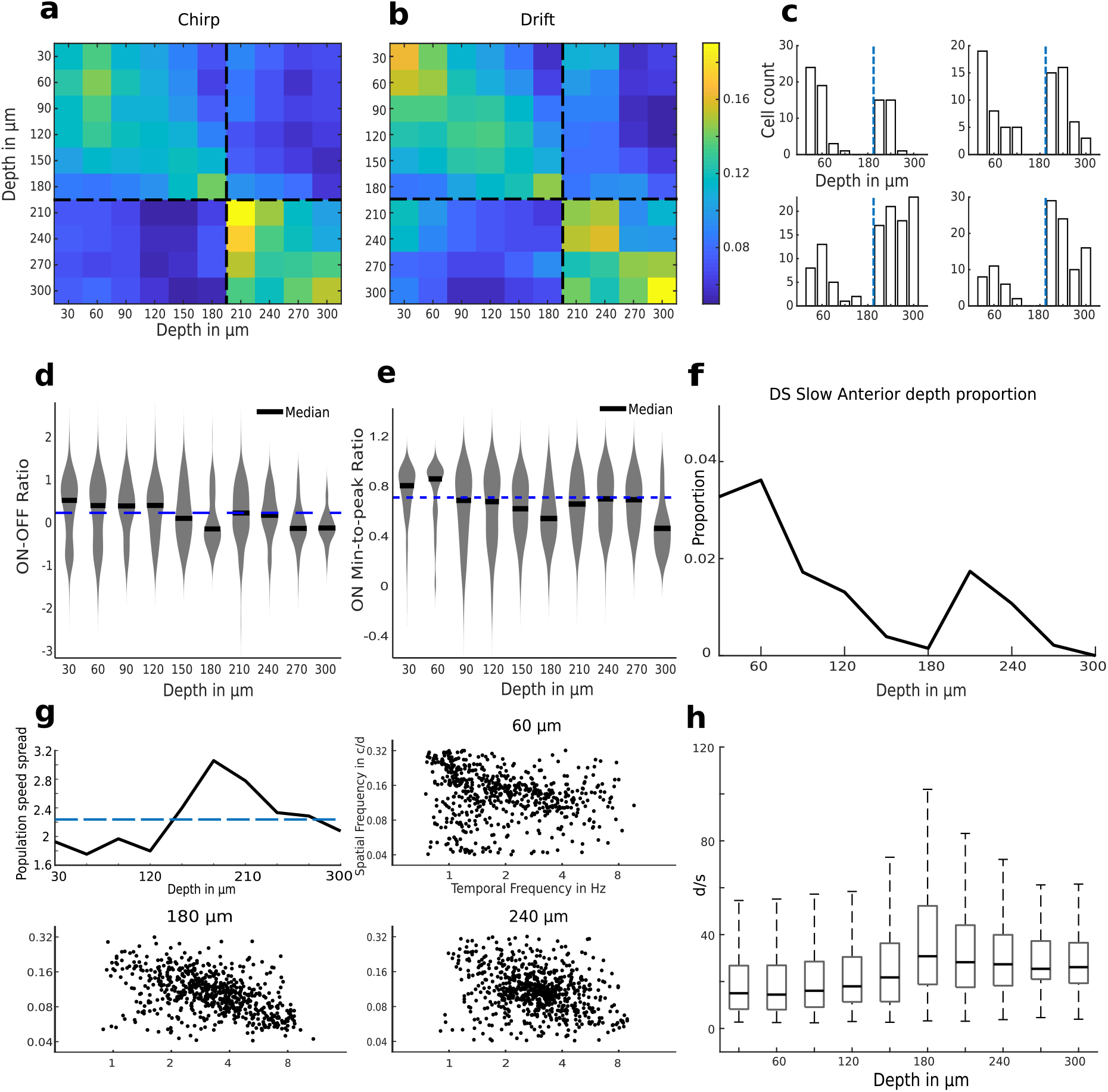
global Depth Changes. **a**) Depth co-clustering correlation matrix for the chirp clustering. Black dashed line indicates the transition between the SGS and the SO. **b)** Depth co-clustering correlation matrix for the chirp clustering presented as in a). **c)** Example histograms of drift (left) and chirp (right) clusters showing co-clustering of superficial SGS and superficial SO cells. Blue dashed line indicates transition between the SGS and the SO. **d)** Violin plots of the On/Off Ratio across depth. Blue dashed line indicates the mean across the whole population. **e)** Same as d) but for On min-to-peak ratio. **f)** Proportion of DS Slow Anterior cells across depth. The proportion falls between superficial (30 µm) and deep (180 µm) SGS (p = 4e-7), increases between deep SGS and superficial (210 µm) SO (p = 0.003), and decreases between superficial and deep (300 µm) SO (p = 0.02). **g)** Top left: Plot of mean population speed spread across depth with mean value shown as a dashed blue line. Population speed spread is higher than mean at 180 µm (p < 0.001). Scatter plots of preferred temporal and spatial frequency for cells at depths of 60 µm (top right), 180 µm (bottom left), and 240 µm (bottom right). **h)** Box plots of speed preference across depth. Preferred speed increases from superficial SGS (30 µm) to deep SGS (180 µm): p = 9e-65. Preferred speed decreases from deep SGS to the SO (200+ µm): p = 2e-4. Both tests were Wilcoxon’s Rank Sum tests.

Having already shown that cardinals of DS subtypes change independently with depth, we inspected cardinal responses throughout the sSC, and observed a tendency for horizontal responses to be represented in the superficial SGS and SO and vertical responses to be represented in the deep SGS. For all DS subtypes, the ratio of vertical to horizontal responses tended to increase with depth in the SGS and then decreased between the deep SGS and the superficial SO, but only significantly for DS Universal cells (p = 5 x 10^−7^ for the SGS increase and p = 0.0053 for the SO decrease). We also observed a double drop in the proportions of anterior motion preferring DS Slow cells throughout the sSC (Figure 5f).

Finally, we plotted the preferred spatial frequency of all cells as a function of their preferred temporal frequency and found a tendency for cells to be located along an orthogonal to the speed isoline (Supplementary Figure 13). We quantified this tendency by dividing the spread of the population in the anti iso-speed direction by the spread in the iso-speed direction to generate the population speed spread at a given depth. This revealed an increase in population speed spread with depth in the SGS, peaking at 180 μm, and then a decline between the deep SGS and the superficial SO (Figure 5g). We additionally found a similar pattern of speed preference with depth (Figure 5h). Thus, we uncovered a relationship between the functional responses of SC neurons and their depth in this brain structure.

## Discussion

### The purpose of multidimensionality in the SC

The current classification of SC cells into a handful of morpho-genetic types presumes a low-dimensional organization of morphological and genetic spaces. If that classification system was to be extended into the functional domain, it would require a low dimensional organization of collicular response space. Here we found that collicular response space is at least multidimensional by demonstrating that neuronal responses to Drift and Chirp stimuli are independent on feature, response type, and clustering levels. A high-dimensional response space would be beneficial to collicular processing in several ways. First, an increase in dimensionality would increase the number of partitions that can be implemented by a downstream linear decoder^41^, resulting in higher computational power. Second, reducing correlations increases efficiency by making the neural code less redundant^2^. Finally, the complex behavior and higher cognitive processes carried out by the SC^7,10,12^ likely require a high-dimensional response space structure^2,5,6^.

Prior RGC classification studies have suggested that Drift and Chirp responses are mutually dependent in the retina^13,32,42^, although this has to be validated experimentally in the future. Based on this assumption, we hypothesize that decoupling must take place at the retinocollicular synapse (Figure 6a). One potential mechanism underlying the decoupling is the functional clustering of local dendritic inputs described at the retinogeniculate synapse^43^. Beyond increasing the dimensionality of response space, decoupling could serve as a preparatory step for general pattern separation or mixed selectivity processes that have been shown in other brain regions^4,44^.

**Figure 6.**
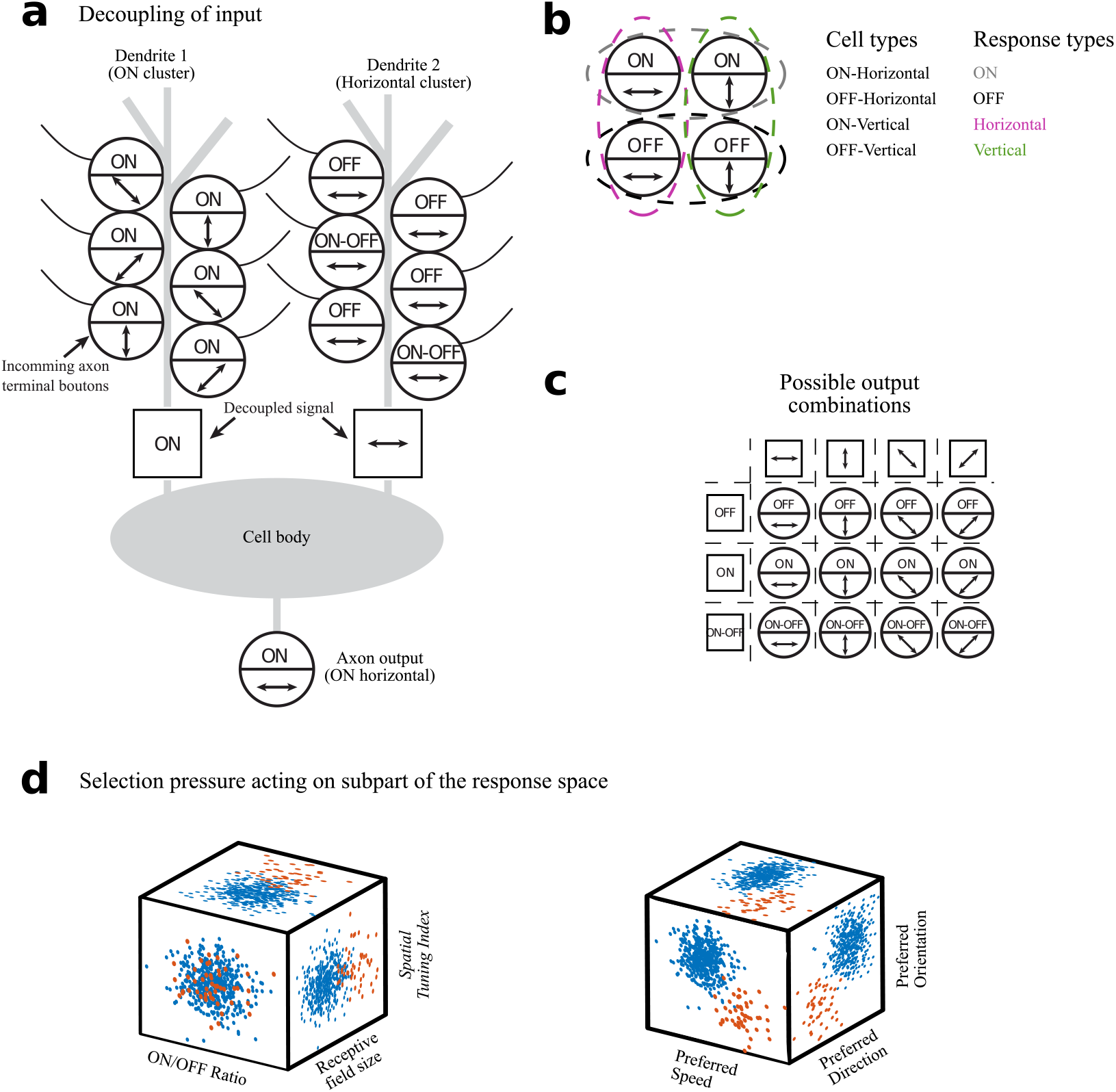
Retino-collicular information transfer. **a**) Model of decoupling of functional responses at the retino-collicular synapse. Such a decoupling would explain why responses that are coupled in the retina are independent in the SC and would allow the generation of new combinations such as ON DS posterior. **b**) Demonstration of the differences between the cell type model and the response type model. Four cells carrying different combinations of responses are depicted on the left. Whereas the cell type model would identify each cell as its own type (middle column), the response type model would identify four responses (right column) with each response type being distributed across several cells (colored dashed lines in the depiction on the left). **c)** Due to the independence of the decoupled responses, all combinations of independent responses will exist. As a result, describing response types is simpler than describing cell types. **d)** Left: Blue and red dots represent the responses of two hypothetical groups of cells. The red group has had their receptive field size changed due to a selection pressure. This pressure did not change the ON/OFF ratio or the Spatial Tuning Index of the group. Right: The selection pressure that caused differences in preferred speed between groups also caused differences in preferred direction and orientation. This in turn lead to higher-than-chance Mutual Information between preferred speed, direction, and orientation. If independencies in response space exist, then selection pressures must be able to act on a subcellular level making response types more tightly aligned with the definition of an evolutionary unit than cell types.

### Classification by response types, not cell types

While the multidimensional structure of collicular response space is problematic for the cell type model, it still has clear structure that merits defining types (Figures 3-4). For this purpose, we propose a response type model where the target for classification is the response itself rather than the cell carrying the response (Figure 6b). The response type model has several advantages over a cell type model. First, it is more concise as it scales additively with each additional independent response whereas the cell type model scales multiplicatively (Figure 6c). We here only tested two responses, but it is possible that many more independencies will be uncovered as more stimuli are tested. Second, if you know that two responses are independent, naming the specific combination of responses in each cell yields no additional information, the additional complexity of the cell type model offers no advantage over the response type model. Third, the response type model allows classification to be used within the framework of the neural network doctrine^45^ by allowing cells to engage in multiple independent pathways. Finally, response types are more in line with the intuitive definition of a type as an “*evolutionary unit with the potential for independent evolutionary change*”^46^ since functional responses directly influence survival whereas functional cell types only influence survival through its impact on the carried functional responses. Together with the capacity of cells for subcellular plasticity^43,47–49^, our finding of response space independencies suggest that the true evolutionary units in collicular response space are response types (Figure 6d).

### Morphology and genetics may not hard-code response types

Our finding of independence among responses predicts that genetic and morphological types cannot have predefined responses to both luminance and motion. Indeed, studies have found a high degree of functional diversity within each genetic subtype in the SC^50,51^. For instance, even the Cbln4-positive cells, which had the highest concentration of direction-selective cells, contained one-third of the cells that were not directionally selective^51^. We hypothesize, that while a cell’s morphology and position in depth are likely determined genetically^20^, a cell-type-specific connectivity with retinal ganglion cells and/or local dendritic clustering of retinal inputs are determined stochastically guided by local plasticity rules^52^. This hypothesis would mean that genes set the stage by increasing the likelihood that a cell will have a particular response type without enforcing a fixed functional role. Given that functional inputs and characteristics vary with depth, it is not surprising that such local stochastic mechanisms lead to marked functional differences among cells from the same genetic type.

### The influence of retinal inputs and anatomical layers

Previous studies have found that collicular direction selectivity is likely inherited from the retina^53,54^. Direct inheritance of direction selectivity is consistent with our finding that collicular DS cells are sensitive to motion in the four cardinal directions detected by the retina (but see^55^). In this respect, our finding that DS subtypes with distinct differences in preferred directions and speed preferences appear to constitute unique response types raises the question of whether a similar subdivision may exist at the retinal level.

We found functional changes with depth in the SC that were common to both Chirp and Drift responses. Specifically, a continuous functional change through the SGS, a steep reset between the SGS and SO, and a second continuous change through the SO. The steep reset leads to a high degree of similarity in depth-related functional organization between the superficial SGS and superficial SO, suggesting that a computational process performed by the SGS is repeated in the SO. A feature showing this pattern of change across depth is the ratio of vertical to horizontal DS responses; cells in the superficial SGS and SO prefers horizontal motion and cells in the deeper SGS and SO prefers vertical motion. This finding aligns well with the finding that, posterior motion-preferring OnOff-DS RGCs project specifically to the superficial SGS^56^, whereas upward motion-preferring Off-DS RGCs project more broadly throughout the SGS^57^. Similar depth-related changes in preferred motion, supported by DS RGC projection patterns, have been described in the dLGN^37,56,58^.

Finally, scatterplots of spatial and temporal preferences in the SC (Supplementary Figure 13) suggest that a selection process pushes collicular cells to respond to a large range of speeds. This tendency is much stronger in the deep SGS than in the superficial SGS and SO, suggesting that speed is more actively selected for in the deep SGS than in other parts of the sSC.

As well as providing insight into the processing of visual information by different layers of the sSC, our findings reveal the limitations of using one-dimensional classification in functionally complex brain structures. As an alternative, we highlight how multidimensional response types can be used to classify high-dimensional response spaces at a functional level.

## Acknowledgements

We thank Zoltan Raics for developing our visual stimulation and imaging system, Bjarke Thomsen, Misugi Yonehara, Celine Thiesen, and Esther Helga for technical assistance. We also thank Hiroki Asari, Akihiro Matsumoto, and Lesley Anson for comments on the manuscript.

## Author contributions

O.S.S. and K.Y. conceived and designed the experiments and analyses. O.S.S. performed all experiments and analyzed all the data. O.S.S. and K.Y. interpreted the data and wrote the paper.

## Methods

### Experimental Methods

#### Animals

All experimental procedures were approved by the Danish National Animal Experiment Committee (License number: 2020-15-0201-00452). Twenty-one 12-to 18 weeks old wild-type mice (C57BL/6J from Janvier Labs) of both sexes were used. The mice were housed in a heat-regulated room in groups of two to four per cage with easily accessible food and water. The mice were kept on a standard 12-hour day/night schedule and looked after daily by animal caretakers. Since the mice were at least 12 weeks old when imaging was performed, cells in both the retina and the SC were fully mature ^59^.

#### Pre– and Post-Procedure Protocol

All procedures were performed in a sterile and aseptic environment. Before procedures, mice were anesthetized with a fentanyl (0.05 mg/kg body weight), midazolam (5.0 mg/kg body weight), and medetomidine (0.5 mg/kg body weight) mixture injected intraperitoneally. Dexamethasone (0.2 mg/kg body weight) were administered subcutaneously to prevent edema during and after surgery. During procedures mouse body temperature was kept stable by using a heating plate and their eyes were protected from dehydration using eye ointment (Viscotears, Novartis).

Post procedure protocol included administration of carprofen (0.08ml subcutaneous) and buprenorphine (0.03ml intramuscular) every 8 hours until the mice no longer showed signs of being in pain for up to 48 hours). Anesthesia was reversed by injecting a mixture of Flumazenil (0.5 mg/kg body weight) and Atipamezole (2.5 mg/kg body weight). The mouse was allowed to slowly wake on the heating plate in its cage.

#### Viral Injection

The mice were anesthetized as described. After the head was shaved, the mouse was held steady by ear bars and the skin was disinfected with 70% ethanol. A sagittal (anterior-posterior) cut in the skin was made 1-2 mm left of the midline until lambda and the injection site was visible. Connective tissue covering the skull was removed and a small burr hole was drilled 0.5-0.7 mm caudal to the interaural line 0.3-0.7 mm left of the midline. Skull debris was removed continuously using absorption spears (SUGI, AgnTho’s) lightly soaked in saline.

Virus (AAV1.Syn.GCaMP6f.WPRE.SV40, 2.13 × 10^13^ vg/ml, Penn Vector Core #100837-AAV1) was injected with a glass pipette through the burr hole with the pipette holder tilted ∼25 degrees posterior to ensure expression reached far enough rostrally. A total of 600 nl was injected across five depths (1.65, 1.5, 1.35, 1.2, and 1.05 mm below skull surface) using as low pressure (Picospritzer III, Parker) as possible over a total of 15 minutes. To avoid backflow, we waited further 5-10 minutes before fully retracting the pipette. The skin was then sutured, and post procedure protocol was followed as described above.

#### Head Plate Surgery

To gain visual access to the SC without removing part of the cortex and thereby potentially altering the response of collicular cells ^60^, we adapted a method designed to expose the collicular surface by displacing the transverse sinus with a silicone plug ^61^. The transverse sinus is assumed to not be essential for healthy brain function, as ablation of the sinus has been shown to cause minimal to no neurological symptoms in humans ^62^. With this method, we gain visual access to approximately 15-25% of the posteromedial surface of the SC.

14-21 days after injection the mouse’s head was shaved and the mouse was placed in an ear bar holder. The shaved skin was disinfected with 70% ethanol and a 1-1.5 cm circular patch of skin, including underlying tissue and a portion of neck muscles was removed to expose the skull above lambda and the injection site. To increase glue binding strength superficial cuts in the skull were then made. A custom-made metal head plate was placed on the skull centered on lambda and attached with glue (Super Glue Power Gel/Flex, Loctite). The function of the head plate was to enable stabilization of the mouse’s head during imaging.

#### Silicone Plug Surgery

For the plug surgery, the anesthetized mouse was placed in a head plate holder. A round cranial window slightly smaller than 4mm centered on the point of the sinus divergence was made using a drill. Skull debris was removed continuously with a suction needle and the exposed brain was kept soaked in sterile phosphate buffered saline (PBS). The dura above the SC was removed and a triangular silicone plug (Kwik-sil, World Precision Instruments) attached to a 4 mm round glass window (0.15 mm thickness, Warner Instruments) was inserted above the SC and slowly pushed forward displacing the left transverse sinus and creating visible access to the SC. The glass plate was then glued to the edge of the skull and the headplate. After the glue had dried, normal post-procedure protocol was followed. Optics remained clear for at least 3 months following plug surgery.

#### In Vivo two Photon Calcium Imaging

A minimum of 7 days after surgery and before performing awake two-photon imaging, the mice were trained to remain calm in the imaging setup by completing four training sessions of increasing duration over a period of four days. During the sessions the mice were placed on the imaging stage with their head fixated to the head plate holder with their body protected by a padded cylindrical cover ^63^. During and after each session the mouse was rewarded with chocolate paste to create positive association and habituation to the imaging area.

Each imaging session lasted 1-2 hours with breaks every 15-25 minutes depending on stimulus length. Animals were kept awake during imaging by offering chocolate paste rewards during breaks.

Imaging was performed using a resonant scanning microscope (VivoScope, Scientifica) controlled by SciScan version 1.2 running 30.9 frames per second at a resolution of 512×512 pixels covering an area of 500×500 um. Dispersion-compensated 940 nm light was provided by a mode-locked Ti:Sapphire laser (MaiTai DeepSee. Spectra-Physics) through a 16× water-immersion objective (Nikon, 0.8 NA). We imaged up to 10 depths per mouse, from 30 μm under the surface, at 30 μm intervals, to ensure no cells were recorded twice, down to a maximum of 300 μm which was the deepest position where we still had adequate signal-to-noise ratio. In our recordings the border between the SGS and the SO was consistently found between 185 and 205 μm below the surface (Supplementary Figure 14). We identified it during imaging by locating the shift in cell body size between lower SGS and SO (Supplementary Figure 14a-c). To determine if the shift in cell body size indeed marked the border between the SGS and SO we imaged Ntsr1-Cre-labeled cells as these cells are known to only reside in the SO ^20^. A quantification of the depth distribution of Ntsr1-Cre cell bodies shows that the emergence of Ntsr1-Cre cells coincides with the shift in average cell body size (Supplementary Figure 14c-d). As such 180 μm was always the deepest SGS layer and 210 μm the shallowest SO layer. For each layer one to three non-overlapping fields of view were imaged.

After fixating the mouse on the imaging stage, the two-photon microscope was moved into place, by putting water on the tip of the microscope and slowly lowering it onto the cranial window glass surface until the surface tension broke, creating a cylinder-shaped water connection between the microscope and the cranial glass window. After this, a layer of black tape was attached covering the open space between the microscope and the metal head plate. This ensured no light contamination from the stimulus screen entered the microscope and slowed down the rate of evaporation of the immersion water. Each imaging session lasted approximately 2 hours, during which the mouse was rewarded with chocolate paste twice, in connection with refilling the water between the microscope objective and the glass window.

#### Screen Positioning

A 47.7 x 26.9 cm 60 Hz screen (Dell, U2212HMc) was positioned 22 cm away from the right eye, angled such that the mean receptive field position of all cells within the field of view was at the center of the screen. Light intensity was measured to 0.051 mW/cm^2^ with a power meter (PM200, Thorlabs). The position of the screen was optimized by testing various screen positions before each imaging session.

#### Stimulus Battery

Mice were presented with the following stimulus battery:

1. **Sparse noise**: Black or white squares covering 10×10 degrees of the visual field were flashed one at a time for 0.1 seconds at all possible xy-positions of a roughly 20-by-10 grid using 5-degree increments in pseudo-random order to determine receptive field position. Grid size was set to the smallest possible size that fully covered the receptive fields of all responsive cells within the field of view.
2. **Drifting Grating (Drift)**: A full-field sinusoidal grating (100% contrast, 0.08 cycles per degree) drifting at a speed of 5 or 40 degrees per second, each presented at 12 equally spaced angles.
3. **Chirp stimulus**: A 10 degree in diameter circular spot was presented in 7-12 different positions on the screen. The chirp stimulus was presented after estimation of receptive fields with the sparse noise stimulus such that the positioning of the chirp circular spot could be optimized to cover the receptive field center of as many cells as possible. The stimulus has three phases. 1. A step phase where the stimulus switched between black and white, 2. A chirp phase where sinusoidal shifts between black and white occurred with frequency increasing in steps, 3. A contrast phase where sinusoidal shifts occur with increasing contrast.
4. **Spatio-Temporal Gratings (SpaTemp):** A full-field sinusoidal grating drifting at all combinations of 8 spatial frequencies (0.04-0.32 cycles per degree, linear increments) and 6 temporal frequencies (0.5-16 Hz, logarithmic increments), each presented at 4 equally spaced angles.

Stimuli were presented in the above-mentioned order, but internally the order of each stimulus was pseudo-random. Between each trial there were 3 seconds without stimuli to allow the cell calcium signals to return to baseline. All stimuli conditions were repeated across 6 trials.

#### Stimulus Size

We chose the size of both the sparse noise (10×10 degree square) and the Chirp stimulus (circle with a 10 degree diameter) based on results from Wang and colleagues ^64^.

## Data Analysis

### Correction of x-, y-, z-axis movement

We manually performed z-axis corrections during imaging by recording a template image before the start of each imaging session and performing re-alignment to the template between each stimulus. Trials with z-movement where at least 5 % of cells showed a simultaneous decrease in fluorescence were discarded.

We performed xy-movement registration by calculating a template using the 20 imaging frames with the highest internal correlation and then using this template to perform correlation-based image registration ^65^. Rigid motion correction was performed with a custom script based on the MATLAB *xcorr* function. We record images at 30.9 frames per second, hence motion across the frame is uniform enough to use rigid plane correction ^65^.

### Neuropil Decontamination and Baseline Calculation

Regions of interest (ROIs) were drawn manually in ImageJ. Neuropil decontamination was performed using FISSA ^66^. For chirp responses, baseline was extracted by running a 0.75 second moving mean, followed by a 60 second 10^th^ percentile filter, and finally another 0.75 second moving mean on the neuropil subtracted fluorescence trace. In the baseline calculation for all other stimuli the 10^th^ percentile filter was replaced with a 20 second moving minimum filter.

Finally, because FISSA in some cases will cause zero values in response amplitude, ΔF/F was calculated by dividing the baseline subtracted response with the median response amplitude of the *pre-neuropil removal* pre-stimulus phase.

### Quality Control

Cells were included if one of the following criteria were met:

1. R^2^ > 0.5 for a 2D gaussian fitted to the response to sparse noise, and Chirp stimulus presented such that it covered the receptive field center of the cell.
2. Chirp Quality Index > 0.45
3. Drifting Quality Index > 0.45, R^2^ > 0.5 for a 2D gaussian fitted to the *pre-neuropil removal response* to sparse noise, and Chirp stimulus presented such that it covered the receptive field center of the cell.

Quality Indexes were calculated as implemented before ^13^. Briefly, if c is the r by t response matrix, r is the number of repetitions of the stimulus, t is the response across time, and 〈〉_x_ and Var[]_x_ denotes the mean and variance across the indicated dimension, respectively, then the quality index (QI) becomes:

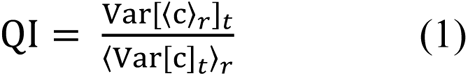

When defined as such a QI can take values between 1/r and 1 with a value of 1 indicating that the exact same signal was recorded across all trials and a value of 1/r indicating no correlation across trials.

### sPCA based Feature Extraction

We used the same method as Baden and colleagues to extract features from Chirp and Drift ^13^. We extracted 20 Chirp features with 10 non-zero time bins (Supplementary Figure 2) by applying sparse principal component analysis (sPCA) ^67^ using elastic net regression as implemented in the SPaSM toolbox for MATLAB ^68^. Before sPCA data was normalized by using the *normalize* function in MATLAB on the mean response across trials.

To extract Drift features, we first performed a singular value decomposition on the normalized mean response across trials. We then used sPCA on the first column (accounting for 86 % of variance) of the resulting temporal components to extract 8 features with 5 non-zero-time bins (Supplementary Figure 3) for both drifting speeds. After extraction the features were z-scored using the *zscore* function in MATLAB.

### Response properties from the Drift Stimulus

We defined a cell as direction-selective (DS) if it had a direction selectivity index (DSi) greater than 0.25 and a permutation test found a p-value of less than 0.05.

The direction selectivity index (DSi) was calculated as defined by Mazurek and colleagues ^69^:

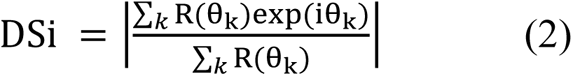

where R(θ_2_) is the maximum response amplitude during the *drifting* phase minus the mean response during the final 0.5 seconds of the *static* phase (Supplementary Figure 1b) for direction θ_k_ (Using directions 0-330 degrees in 30 degree intervals).

A cell was defined as DS Slow if it had a DS response to the slow grating but not to the fast grating. Similarly, a cell was defined as DS Fast if it had a DS response to the fast grating but not to the slow grating. Finally, a cell was defined as DS Universal if it had a DS response at both speeds. As such, DS Slow, DS Fast, and DS Universal are mutually exclusive.

We defined a cell as orientation-selective (OS) if it had an orientation selectivity index (OSi) greater than 0.25 and a permutation test found a p-value of less than 0.05.

The orientation selectivity index (OSi) was calculated as defined by Mazurek and colleagues ^69^:

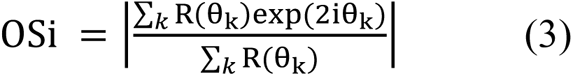

where R(θ_2_) is the maximum response amplitude during the *static* phase minus the mean response during the final 0.5 seconds of the *pre-stimulus* phase (Supplementary Figure 1b) for direction θk (Using directions 0-330 degrees at 30-degree intervals).

For both orientation– and direction selectivity the preferred orientation/direction was calculated as the angle of the summed vector.

Permutation tests were done by shuffling responses across all trials and directions 1000 times, calculating the DSi/OSi of each shuffle, and performing t-tests on the actual DSi/OSi vs the DSi/OSi of the shuffled responses.

### Response properties from the Chirp Stimulus

We used the first six seconds of the mean response across trials to the Chirp stimulus to define On/Off subtypes. The first two seconds of the response were used to quantify baseline response and standard deviation, the following two seconds quantified the ON response, and the last two seconds quantified the OFF response (Supplementary Figure 1c-d).

We define *ON response* as the maximum response amplitude during ON phase minus the mean response during the last 0.5 second of pre-stimulus phase, and *OFF response* as the maximum response amplitude during OFF phase minus the mean response amplitude during the last 0.5 second of the ON phase (Supplementary Figure 1c-d).

The ON-OFF ratio for a cell was defined as follows:

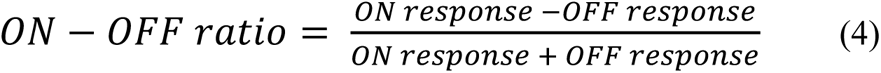

A cell was defined as an On cell if the ON response was at least 2.75 standard deviations above baseline and the OFF response was at most 1.5 standard deviations above baseline. A cell was defined as an Off cell if the OFF response was at least 2.75 standard deviations above baseline and the ON response was at most 1.5 standard deviations above baseline. Finally, a cell was defined as an OnOff cell if both the ON and the OFF response were at least 2.75 standard deviations above baseline. The 2.75 standard deviation threshold was chosen because it was the point at which the chance of getting a false positive was 1 % when using normally distributed random data under the same conditions. The 1.5 standard deviation cutoff was set after testing a wide variety of values and manually inspecting responses of the resulting On and Off cells. Specifically, using a higher cutoff value caused the mean response of the Off population to have visible ON responses and the mean response of the On population to have visible OFF responses (Supplementary Figure 15). All statistical tests based on definitions of On and Off cells were run using both a 1.5 and a 2.75 cutoff. We found no differences in the results of the tests between the two conditions.

We further extracted two parameters that quantified the dynamics of the ON and OFF response:

1. **time-to-peak** measures how long time it takes from the onset of the stimulus until at least 90 % of peak amplitude is reached (Supplementary Figure 1d).
2. **min-to-peak ratio** measures how much of peak amplitude was lost 0.5 seconds following the peak (Supplementary Figure 1d) and was calculated as follows for On, Off, and OnOff ON responses:

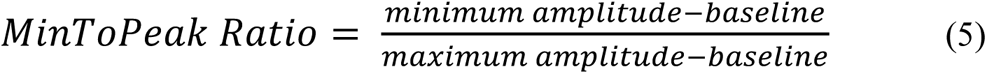

where *maximum amplitude* is the maximum amplitude during the given phase, *minimum amplitude* is the minimum amplitude during the 0.5 seconds following the *maximum amplitude*, and *baseline* is the mean response during the last 0.5 seconds before the onset of the given phase. Min-to-peak ratio for OnOff OFF responses were calculated using the same method but the baseline used were the mean response during the last 0.5 seconds of the pre-stimulus period to avoid loss of amplitude of sustained ON responses to cause the min-to-peak ratio to get high negative values (Supplementary Figure 1d).

On/Off responses have often been divided into Fast/Slow and Transient/Sustained based on response dynamics ^13,32^. To do the same, we performed k-means clustering with the number of clusters set to two, on the min-to-peak ratio and time-to-peak data for On, Off, and OnOff cells (Supplementary Figure 16). The resulting segregation of Sustained and Transient responses is supported by visual inspection of the histograms (Supplementary Figure 16a). The k-means clustering processes were restarted 50 times and the stability of the resulting clusters were assessed. In all cases the threshold changed less than 0.01 between restarts. The application of k-means to the time-to-peak data did not align with the visual data, as visual inspection of the histograms indicated a need for a different, potentially a further, subdivision (Supplementary Figure 16b). Furthermore, the thresholds set by clustering showed considerable changes across restarts (shifts in threshold of up to 0.26 seconds). On this basis we manually defined a further border based on the visual inspection (Supplementary Figure 16c) and plotted depth distributions of the three subdivisions (Fast, Medium, Slow) for all four On/Off responses (Supplementary Figure 16d). Since the Medium group was more similar in depth distribution to the Slow group for On, OnOff-ON, and OnOff-OFF, we chose to place the Slow/Fast division along the Medium/Fast border rather than at the Medium/Slow border.

Statistical significance of differences in ON/OFF response properties between DS and non-DS cells were calculated using a Wilcoxon’s rank sum test corrected for multiple comparisons by Bonferoni-Holm correction and Wilson’s Harmonic Mean.

### Features from the SpaTemp Stimulus

Responses to the SpaTemp stimulus were first averaged across trials and directions. Then the response amplitude for each spatio-temporal frequency combination was calculated by extracting the max response during the drifting phase and subtracting the mean response during the pre-stim phase (Supplementary Figure 1e) resulting in an x by y matrix of response amplitudes for each cell where x and y are the number of temporal and spatial frequencies respectively. Five features were extracted from this matrix:

1. **Temporal Tuning** is a measure of how selective a cell is with respect to temporal frequencies^70^. It was extracted by first calculating the mean of the response amplitude matrix across spatial frequencies, resulting in a 1 by x matrix for each cell where x is the number of temporal frequencies (from here on out called the *temporal response curve*). Then for each cell we found the maximum response amplitude of the temporal response curve and divided it by the sum of the curve.
2. **Spatial Tuning** is a measure of how selective a cell is with respect to spatial frequencies. It was extracted in the same way as temporal tuning with the exception that because we measured spatial tuning at linear intervals, we first converted the spatial frequencies to log intervals by subsampling.
3. **Temporal preference** is a measure of a cell’s preferred temporal frequency. It was calculated by extracting the temporal response curve, up-sampling it by a factor of 1000, and finding the equal-area point of the up-sampled curve. To reduce the influence of noise a small constant was added to the temporal response curve before up sampling. This forces unresponsive cells to have a temporal preference of around 1 divided by the number of temporal frequencies instead of having a random temporal frequency decided by noise. Forcing unresponsive cells to have similar values is desirable for features used for clustering as it causes unresponsive cells to end up in the same cluster instead of being spread out across all clusters randomly. The value of the added constant was set to 0.001 dF/F as this was the median response amplitude across all conditions of unresponsive cells.
4. **Spatial preference** is a measure of a cell’s preferred spatial frequency. It was calculated in the same way as temporal preference.
5. **Speed preference** is a measure of a cell’s preferred speed. It was calculated by taking a cell’s preferred temporal frequency and dividing it by its preferred spatial frequency.

### Functional Clustering

We used Gaussian Mixture Modeling (GMM) ^71^ for our cluster analyses with Bayesian Information Criterion (BIC) ^72^ as a penalizing term to avoid overfitting when choosing the optimal number of clusters. The BIC was calculated as:

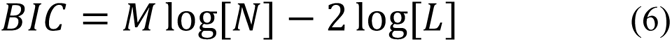

where M is the number of parameters estimated by the model, N is the number of cells, and L is the likelihood of the model.

Our full clustering procedure consisted of the following five steps (Supplementary Figure 9A-C):

1. The extracted features were fitted with a GMM (as implemented in the MATLAB *fitgmdist* function) ten times with predefined numbers of clusters ranging from 1 to 100. The BIC was then evaluated for each number of clusters across the ten repeats, and the number of clusters that yielded the lowest BIC was selected.
2. Using the model selected by the BIC evaluation we performed a further 30 GMM clusterings and extracted a co-clustering matrix (an n by n matrix where point (x,y) lists how often cell x and cell y ended up in the same cluster across the 30 repeats).
3. We then sorted the co-clustering matrix by performing agglomerative hierarchical clustering, as implemented in the MATLAB *linkage* function, on the matrix and used a custom-made script to define new clusters based on their co-clustering score. The script first reduced the matrix by excluding loosely clustered cells defined as cells that end up in the same cluster as their 10 most similar cells less than ⅔ of the time. Then cluster edges were detected in the reduced co-clustering matrix and used to define the borders of the new co-clustering score-based clusters. Finally, the loosely clustered cells were re-introduced by adding them to the cluster with which they had the highest mean co-clustering score.
4. We then generated 25 random subsamples each containing 90 % of the cells from the original dataset and repeated steps 1-3 to generate 25 subsample clusterings and calculated the Jaccard similarity score (see section 2.2.4.3) between each cluster in the original clustering and the most similar cluster in each of the 25 subsamples. Clusters with a mean Jaccard similarity score of less than 0.5 across the subsamples were discarded and their cells re-assigned to the cluster with which the merging cell had the highest average co-clustering score.
5. Finally, dendrograms were created by using agglomerative hierarchical clustering (as implemented in the MATLAB *linkage* function) on the mean feature values of each cluster with distance defined as average unweighted distance.

Steps 1, 4, and 5 are commonly used in functional clustering studies ^13,22^. In addition to this “standard” method we added steps 2 and 3 because we noticed that, even though individual clusters changed between clusterings, the same “meta-structure” was present across all clusterings (e.g., OS/DS subtypes and On/Off subtypes were always placed on separate branches). A key reason for running these clusterings was that we wanted to compare Drift and Chirp clusterings to see if there was a larger than chance overlap between the clusters defined by the two feature sets. Such an overlap could easily be overlooked if repeat-to-repeat variability is high so we chose to focus on the cross-repeat meta-structure to maximize the likelihood that we would find any existing overlap between Chirp and Drift clusters.

To ensure that adding steps that emphasize the meta-structure of the data did not generate non-existing structure by overfitting the data, we tested it on normally distributed random noise to see if our method would generate clusters in random data. However, the GMM followed by BIC consistently (and correctly) estimated the number of clusters to one. Therefore, we tested the model again, but this time ignored the BIC by forcing the GMM to fit the same number of clusters as our real data. Following steps 2-5 we ended with a model that identified four “clusters” in the random data that all had a lower Jaccard score than the 0.5 cutoff we set for the real data (Supplementary Figure 17).

### Batch effect analysis

Whenever a functional imaging study uses data collected across multiple sessions there is a risk of batch effects causing functional responses of cells to differ between sessions. In our case, data was collected across 83 sessions where each session consisted of recording responses from all cells within a single field of view. Because our imaging plane is oriented in parallel to the surface of the SC cells recorded within the same session are from the same depth within the SC. Given that the SC is a layered structure with clear functional changes between layers this would cause you to expect functional differences between sessions.

To test for batch effects, we therefore developed a method that can differentiate between functional differences stemming from batch effects from ones stemming from depth differences. We did this by calculating the session proportions for each cluster as follows:

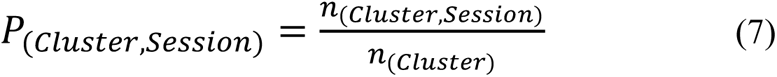

where n_(cluster, session)_ is the number of cells in the cluster stemming from a given session and n_(cluster)_ is the total number of cells in the cluster. To assess whether a cluster’s session proportions deviate from chance, we did a permutation test where the cluster tags were randomized within depth (i.e. cluster tags of cells with their cell body placed at a certain depth below the SC surface were permuted separate from other cells at other depths). For each of these permutations we calculated the session proportions as stated in equation (7) and extracted the expected session proportions for a cluster with the given depth composition by calculating the mean session proportions across permutations for each cluster. We then used the expected session proportions to calculate a deviation from expectation for each cluster as follows:

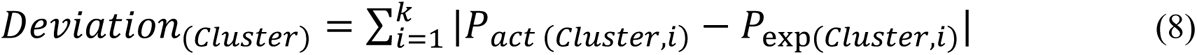

where *P_act_*_(*cluster, i*)_ is the actual proportion of cells from the given cluster that stemmed from session *i* and *P*_exp(*cluster, i*)_ is the expected proportion. Finally, we tested the significance of the deviation by calculating the deviation from the expected session proportions of each random permutation and performing a t-test of these values against the actual deviation.

Using this method, we found that 13 of 50 drift clusters and 14 of 28 chirp clusters deviated more than chance from the expected session proportion (Supplementary Figure 18A-B) possibly indicating batch effects.

Looking across all clusters the Drift clustering deviated on average by 0.8 percentage points from the expected session proportion whereas the random permutations deviated by 0.65 percentage points. For the Chirp clustering the numbers were 0.68 percentage points and 0.51 percentage points, respectively.

Assessing whether these numbers are concerning in relation to the conclusions of our study is difficult for at least three reasons. First, even though a higher-than-chance deviation could stem from influences that should be irrelevant to the functional classification of a cell (such as time of day of the recording, or age and sex of the mouse), it could also stem from differences that are relevant to functional classification such as position of the cell in the visual field as this type of difference would also be correlated with session number. Second, we have not been able to find data on whether this amount of potential batch effect is higher than other similar studies as these do not report considerations regarding batch effects ^13,21,22^. Third, even if a batch effect is biasing our results, it is not given it would affect our main findings as these are based primarily on comparisons between On/Off and OS/DS responses which were both recorded within all sessions and as such should be affected equally by batch effects.

While avoiding batch effects entirely would be preferable it is difficult to implement in large scale imaging studies that require collecting data across multiple sessions. Given that other similar studies are not reporting taking steps to address bias stemming from batch effects this could mean that going into more detail with these effects could lead to important improvements of functional clustering studies. Given more time it would have been beneficial to do a more thorough test to try to identify the source of this potential bias by controlling for things such as xy-position of the FoV, time of day of imaging, age and sex of mouse, mental state of mouse (anxious or relaxed etc.), amount of subdural bleeding during surgery, etc.

## Calculations and statistical tests

### Controlling for Depth

The SC is known to be a highly layered structure. All features we measured had clear changes in values across depth. To avoid these depth changes biasing our tests, all permutations tests were done such that the depth distribution of cells in the random permutations matched the depth distribution in the tested subpopulation.

### Corrections for Multiple Comparisons

To counteract the problem of multiple comparisons we used the Bonferroni-Holm method ^73^, and then, to further increase statistical power, also calculated Wilsons harmonic mean ^74^.

### Mutual information

One way to evaluate the degree of dependency between variables is to calculate their mutual information (MI) ^33^. MI can be defined as follows:

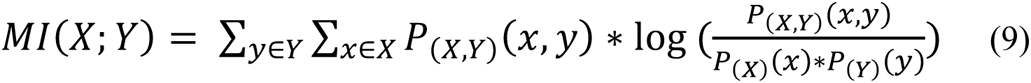

where P_(*X,Y*)_ is the joint probability function and P_(*X*)_ and P_(*Y*)_ are the marginal probability functions^75^. This means that if the joint probability is equal to the product of the marginal probabilities MI will be zero. In other words, if MI between two features is zero or at chance level the features are independent, and consequently any overlap between subpopulations based on those features will be coincidental.

MI was calculated in MATLAB using the *mi_cont_cont* function ^76^. To test for significance of the MI between two features, A and B, we made 500 random permutations of B, calculated MI between A and B and A and the random permutations, and calculated the p-value as the proportion of permutations with higher MI than the actual MI. This calculation is “saturated” at a p value of 1 divided by the number of permutations. In those cases, we increased the number of permutations until the test no longer saturated.

### Proportions and expected proportions

The proportion of a given group of cells at a given depth was calculated as:

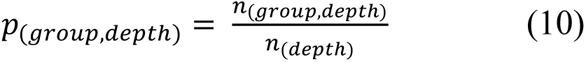

Where n_(group, depth)_ is the number of cells belonging to the given group at the given depth and n_(depth)_ is the total number of cells at that depth.

The expected proportion of a given combination of groups at a given depth was calculated as:

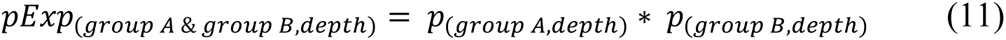

The actual proportion of a given combination of groups at a given depth was calculated as in (10) with *n*_(*group, depth*)_ defined as the number of cells belonging to both groups at the given depth.

The summed deviation between actual and expected proportions was calculated as:

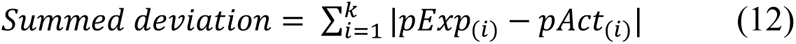

where *pExp*_(*i*)_ is the expected proportion at depth *i*, *pAct*_(*i*)_ is the actual proportion at depth *i*, and *k* is the number of depths. Tests for statistical significance was carried out by permuting group A and group B tags relative to each other within each depth, calculating the summed deviation between each permutation against the expected proportion, and running a t-test of the actual summed deviation vs the summed deviations from the random permutations.

The proportion of a given response type in each cluster was calculated as:

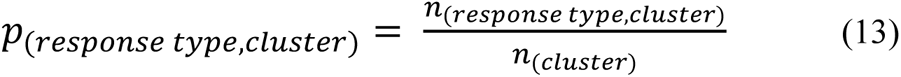

Where n_(response type, cluster)_ is the number of cells in the cluster belonging to the given response type and *n*_(/345067)_ is the total number of cells in the cluster.

The difference between On-DS Universal and OnOff-DS Universal cluster proportions were quantified as follows:

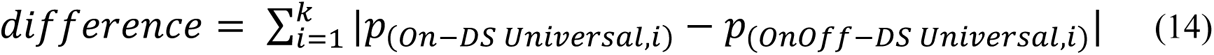

where *p*_(*on-DS universal, i*)_ is the proportion of On-DS Universal cells in cluster *i*, *p*_(_*_onOff-DS universal, i_*_)_ is the proportion of OnOff-DS Universal cells in cluster *i*, and *k* is the number of clusters in the Chirp-Drift clustering. The significance of the difference was estimated by calculating the summed difference in cluster proportions between 500 random samples of On cells of the same size and depth distribution as On-DS Universal cells vs 500 random samples of On cells of the same size and depth distribution as OnOff-DS Universal cells and running a t-test.

### Jaccard scores

When evaluating the stability of our clusterings and when comparing the similarity of different clusterings we used Jaccard Similarity Score ^77^. The Jaccard similarity score between two clusters, A and B, is defined as:

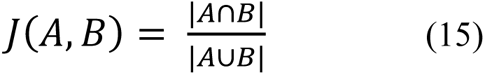

where |A∩B| and |A∪B| denotes the number of cells belonging to both clusters and the number of cells belonging to either cluster, respectively. When used in the context of clustering, it quantifies how similar two clusters are on a scale from 0 to 1 where a value of 1 indicates that the clusters contain the exact same cells ^78^. The significance of the similarity between two clusterings was calculated by permuting the cluster tags relative to each other, calculating the median Jaccard score of the permuted clusters and doing a t-test of the permuted scores vs the actual score.

We wanted to evaluate whether adding orthogonal features or noise to a single feature set would disrupt the clustering more. This was done by comparing the Jaccard score between the chirp clustering and the chirp-drift clustering to the Jaccard scores between the chirp clustering and a number of chirp-noise clusterings. The significance between the difference in Jaccard score was calculated using a t-test. The noise was generated by randomly permuting the n-by-y matrix where n is the number of cells and y is the number of drift features.

### Clustering ability to separate response types

A clustering’s ability to separate response types was quantified in an analogous manner to summed deviation (12) with the addition of being weighted by 1 over the number of clusters to get the mean separation per cluster.

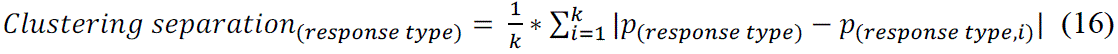

where *p*_(*response type*)_ is the proportion of cells with the given response type across the full dataset, *p*_(*response type, i*)_ is the proportion of cells with the given response type within cluster *i*, and *k* is the number of clusters in the clustering. Significance of the clustering separations was tested by t-testing against 500 random permutations.

### Circular statistics

DS cell cardinal directions were found by fitting a von Mises mixture model ^79^ with four components to each of the three circular histograms shown in 7b and assigning a cell to a component if it was within 22.5 degrees of the center.

Comparisons of preferred direction profiles were carried out by using a two-sample Kuiper test as implemented in *circ_kuipertest* from the Circular Statistics Toolbox for MATLAB ^80^.

### Depth co-clustering correlation matrix

The quantification of how often cells from depths A and B end up in the same cluster was done as follows. First, we counted the number of cells from each depth for each cluster. Since there were differences in how many cells that were recorded from each depth, we corrected for this by dividing the number of cells at each depth with the proportion of cells recorded at that depth across all clusters and rounding the results. We then quantified the number of “connections” between depths by counting across all clusters how many times a cell from depth A appeared in the same cluster as a cell from depth B and normalized the outcome so each row in the matrix sums to one. Since this quantification scales with the number of cells in the cluster squared, we normalized the cluster size before calculating the number of connections in order to avoid large clusters having a disproportionately heavy influence on the outcome.

### Population speed spread

Population speed spread was calculated as follows:

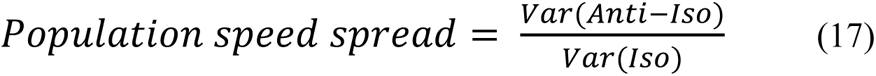

where *Var*(*Iso*) denotes the variance of the population after projecting it onto the iso-speed line and *Var*(*Anti* − *Iso*) denotes the variance of the population after projecting it into a line orthogonal to the iso-speed line.

