## Supplementary Figures for "The superior colliculus response space has globally high– and locally low-dimensionality"

#### *Supplementary figure 1 Stimuli*

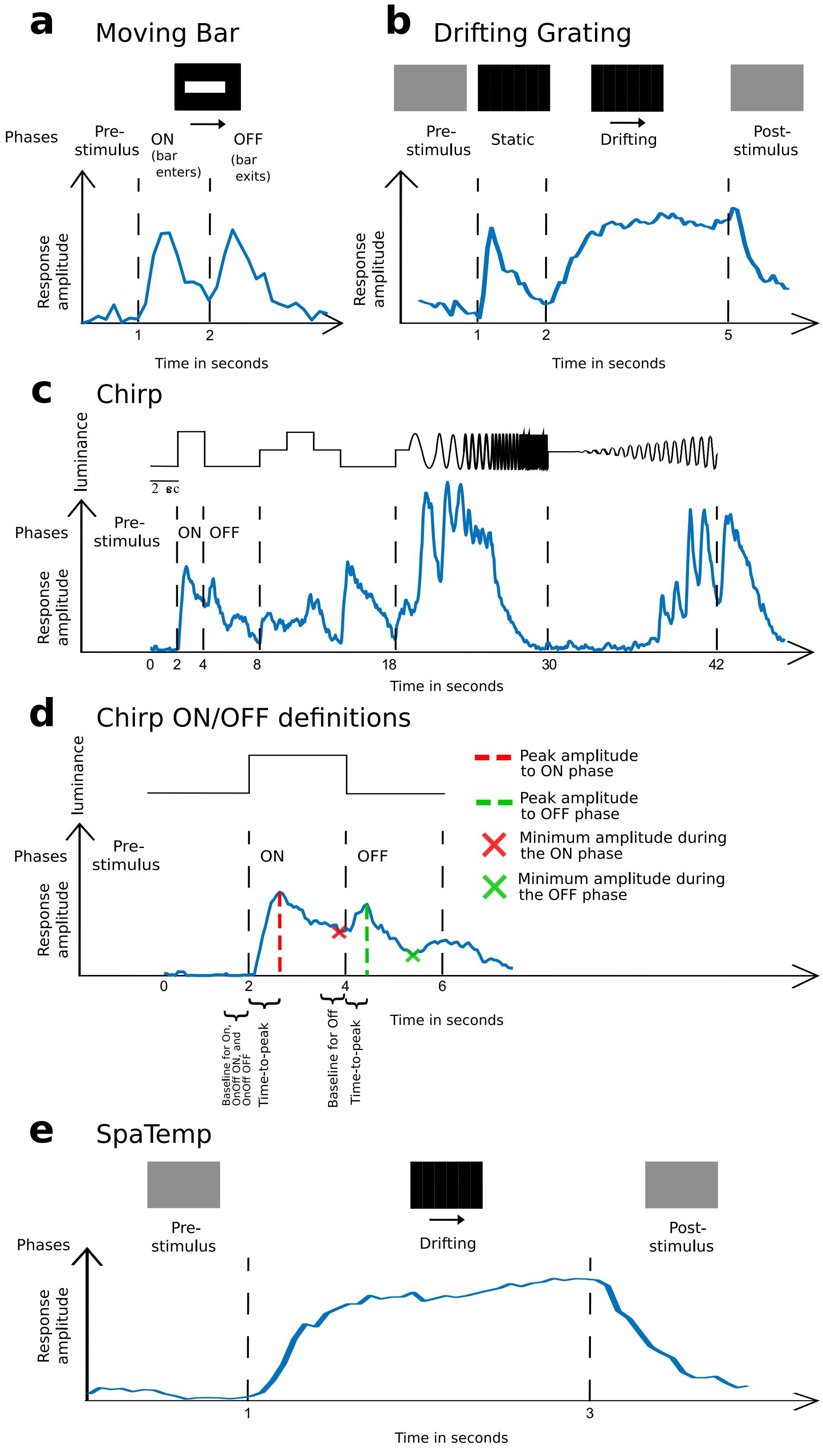


**a)** Moving Bar stimulus from Baden and colleagues with data from the same study[13]: a white bar moves in the longitudinal direction across a black background. Stimuli are presented in 8 equally spaced directions. **b)** Drifting Grating: full-field sinusoidal grating drifting at 5 or 40 degrees per second, in 12 equally spaced angles. **c)** Chirp: a circle, 10 degrees in diameter, alternating in luminance, first with step changes for detecting ON/OFF responses, followed by a chirp phase with sinusoidal shifts in luminance with increasing frequency, ending with a contrast phase with sinusoidal shifts with increasing contrast. **d)** Zoom in of the first ON/OFF step phase of the Chirp stimulus demonstrating how time-to-peak and min-to-peak ratio were calculated. **e)** Timeline of the SpaTemp stimulus.

### *Supplementary figure 2 Chirp Feature Demonstration*

**a**

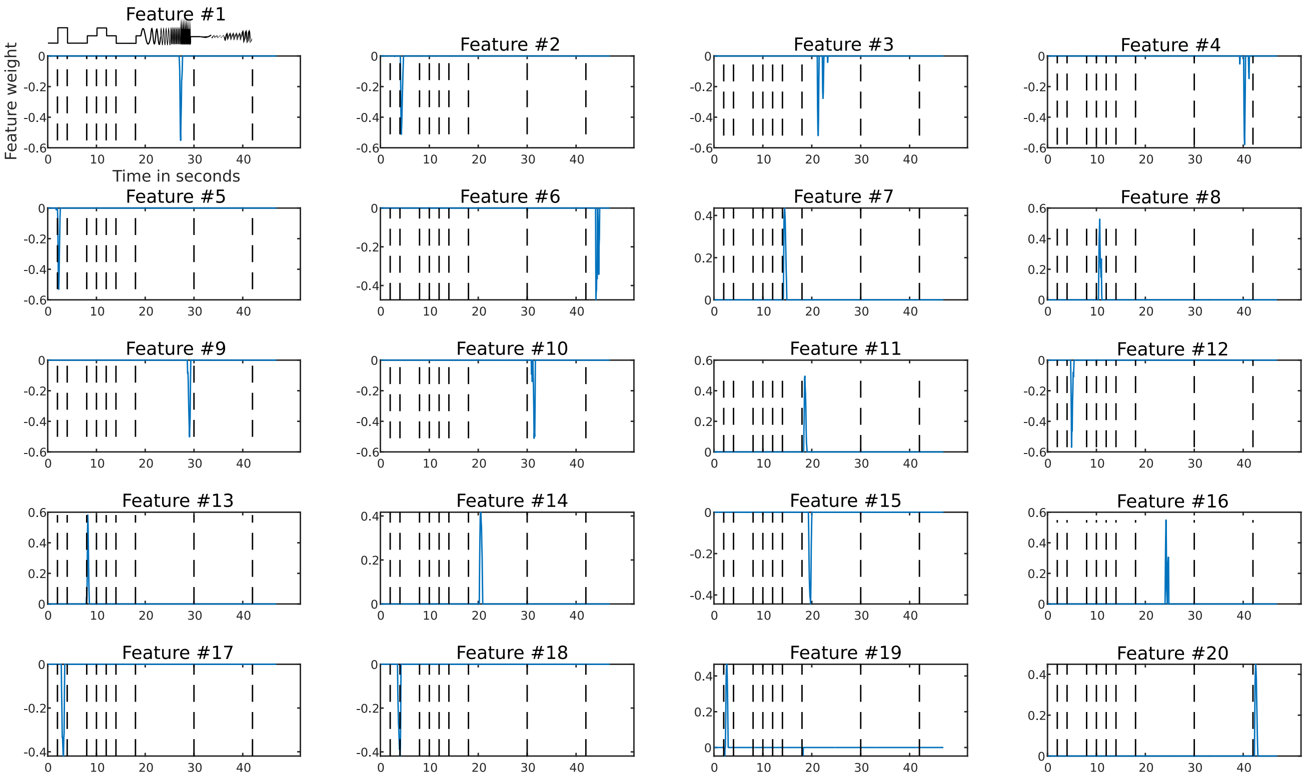


**b**

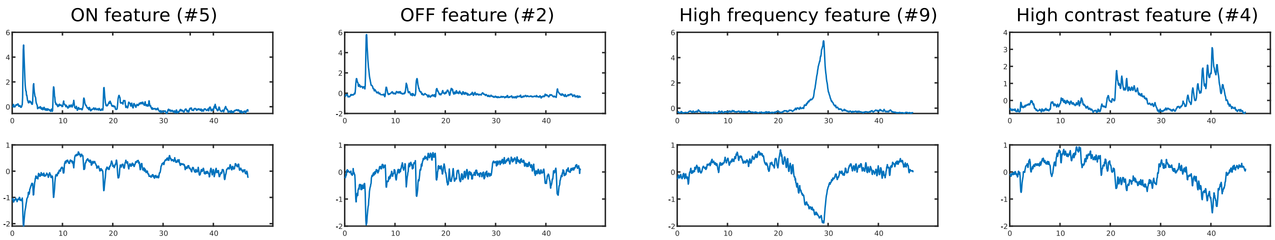


**a)** Plots of each Chirp feature showing the weight as a function of time. **b)** Mean response of the 50 cells with the highest (top) and lowest (bottom) feature score from four example features. Features numbers are written in the title.

#### *Supplementary figure 3 Drift Feature Demonstration*

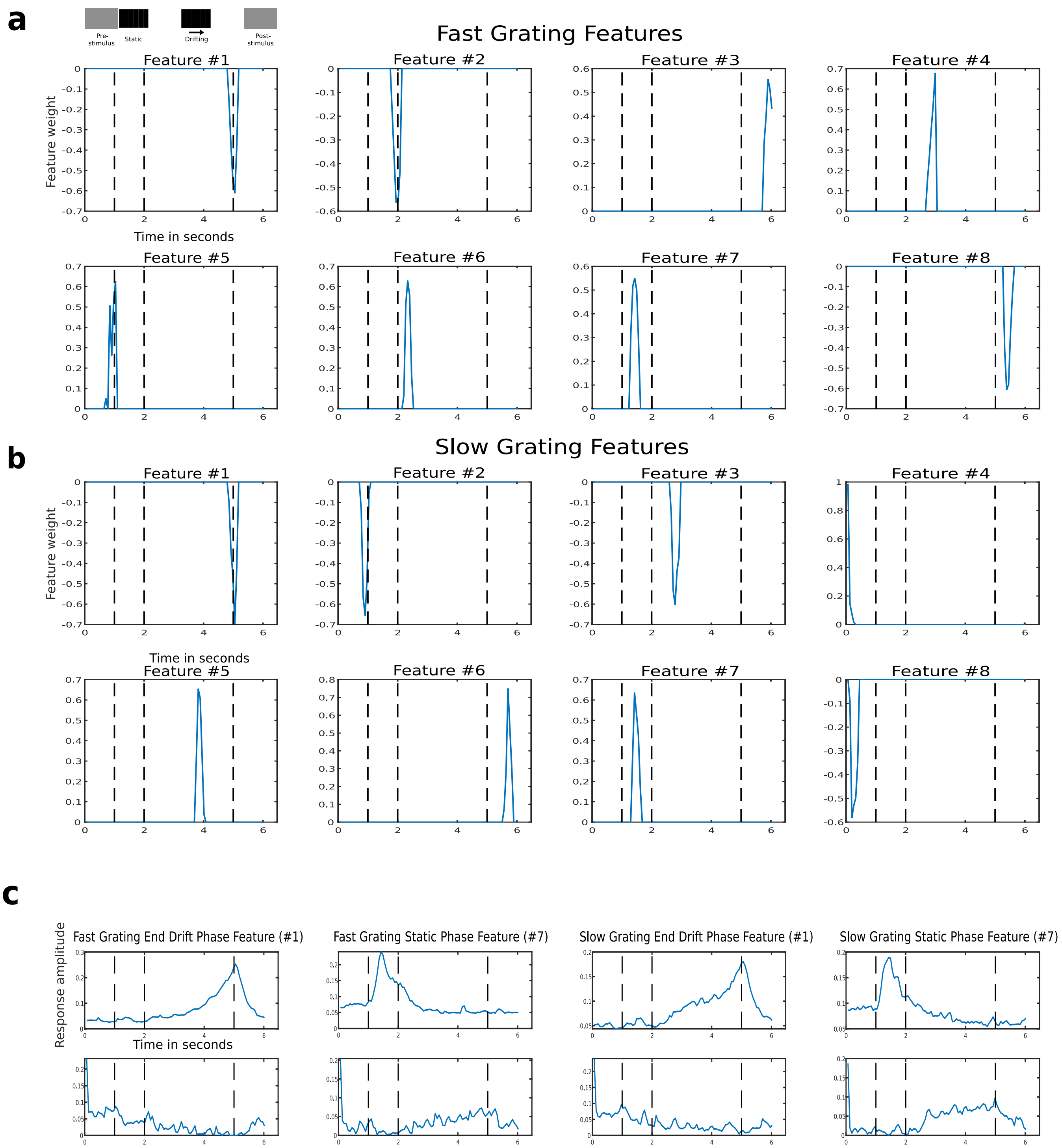


**a)** Plots of each feature from the fast grating stimulus showing the weight as a function of time. Vertical dashed lines indicate transitions between phases of the stimulus as indicated above Feature #1 **b)** Plots of each feature from the slow grating stimulus showing the weight as a function of time. **c)** Mean response of the 50 cells with the highest (top) or lowest (bottom) feature score from four example features.

#### *Supplementary figure 4 Mutual Information between Chirp, Drift, and SpaTemp features*

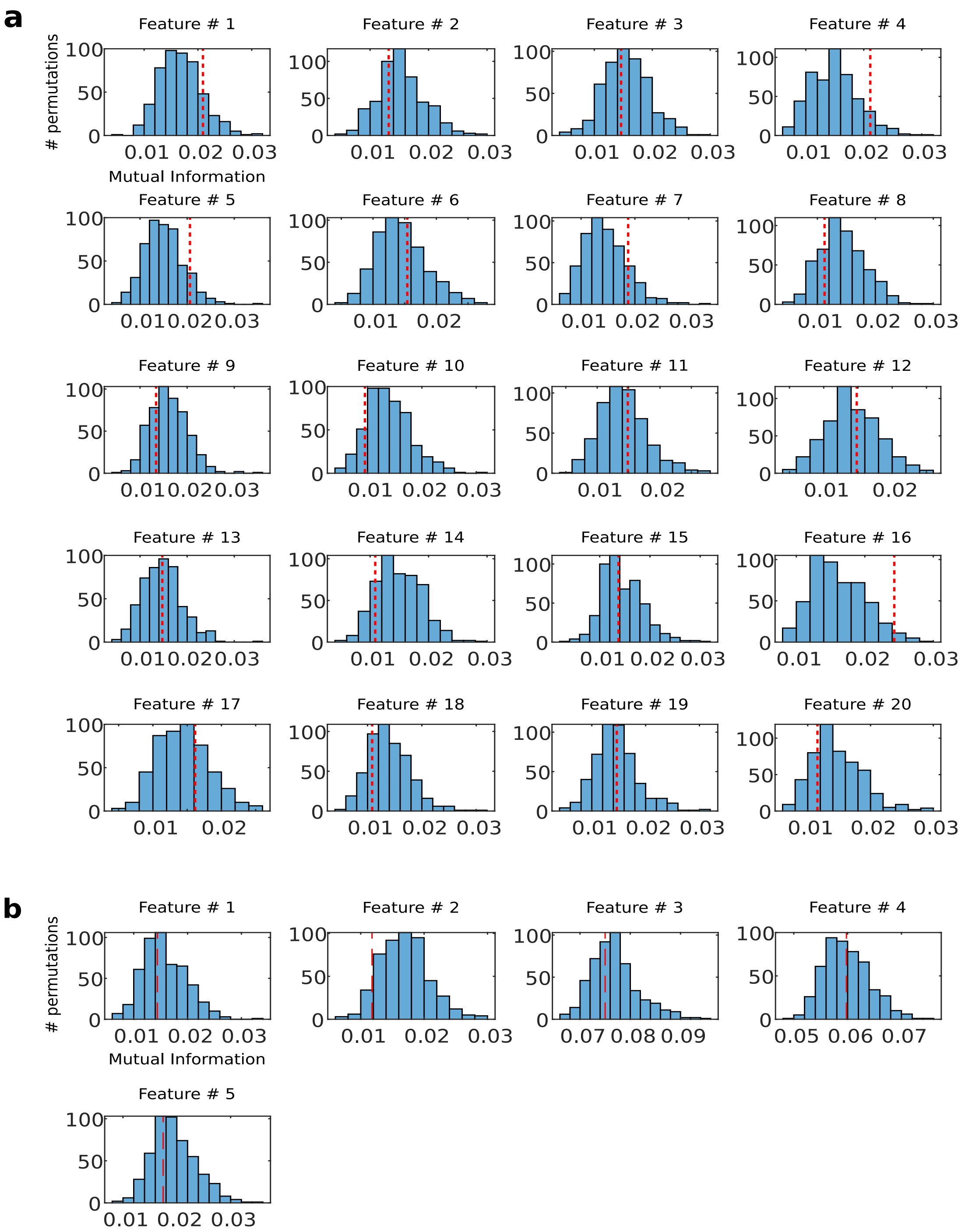


**a)** MI for each Chirp feature vs the Drift feature with which they had the highest MI (dashed red lines) plotted on top of histograms of 500 calculations of MI after the order of the Drift feature had been randomized relative to the order of the Chirp feature. **b)** Same as a) but for MI between SpaTemp features and the Chirp feature with which they had the highest MI.

#### *Supplementary figure 5A Mean Responses of Defined Populations*

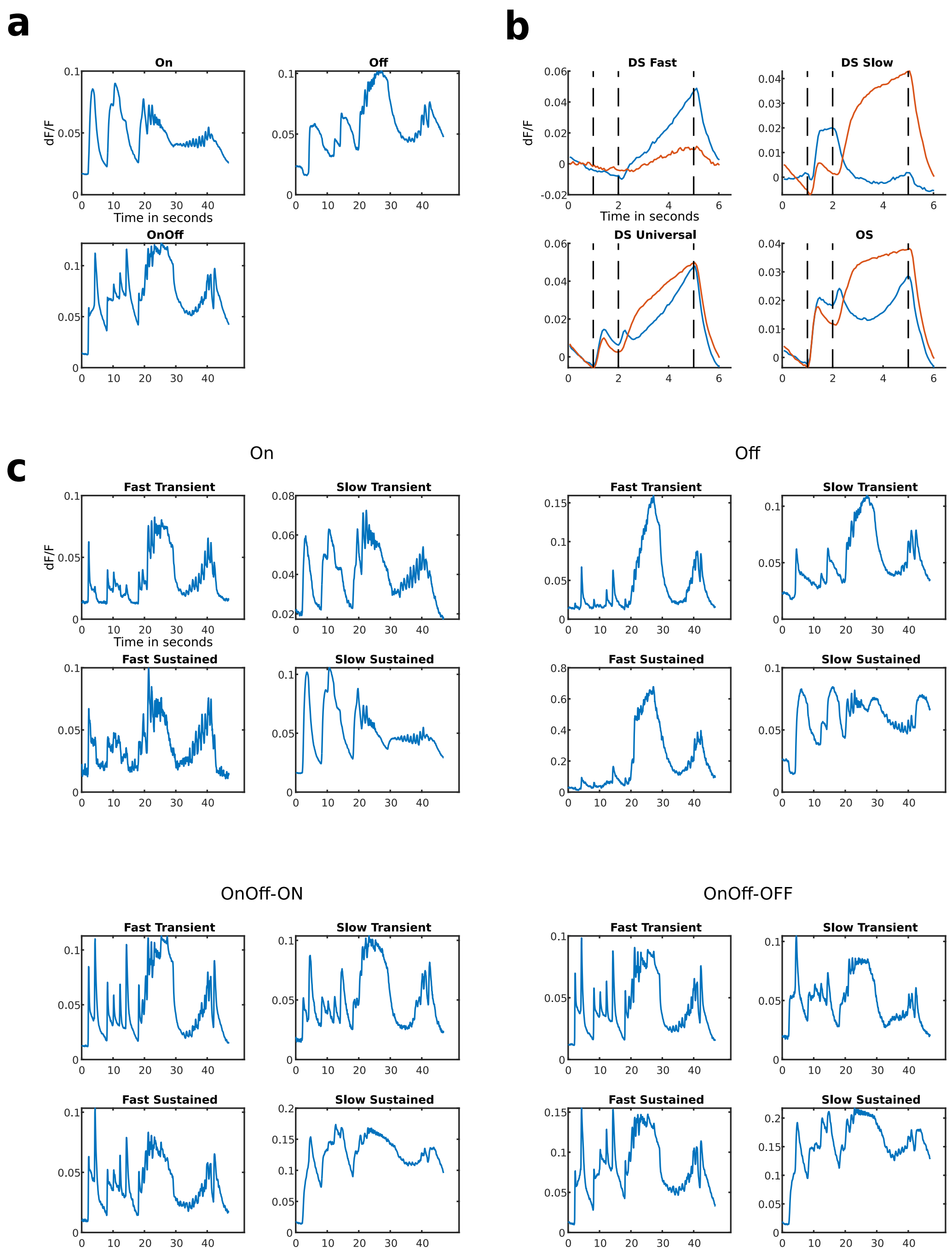


**a)** Mean chirp response of the three On/Off populations. **b)** Mean amplitude of the first component of a singular value decomposition (SVD) of the drift response of each of the four OS/DS populations. The fast grating response is shown in blue and the slow grating response is shown in red. **c)** Mean chirp response of each of the 16 On/Off subpopulations.

#### *Supplementary figure 5B Example Responses of Defined Populations*

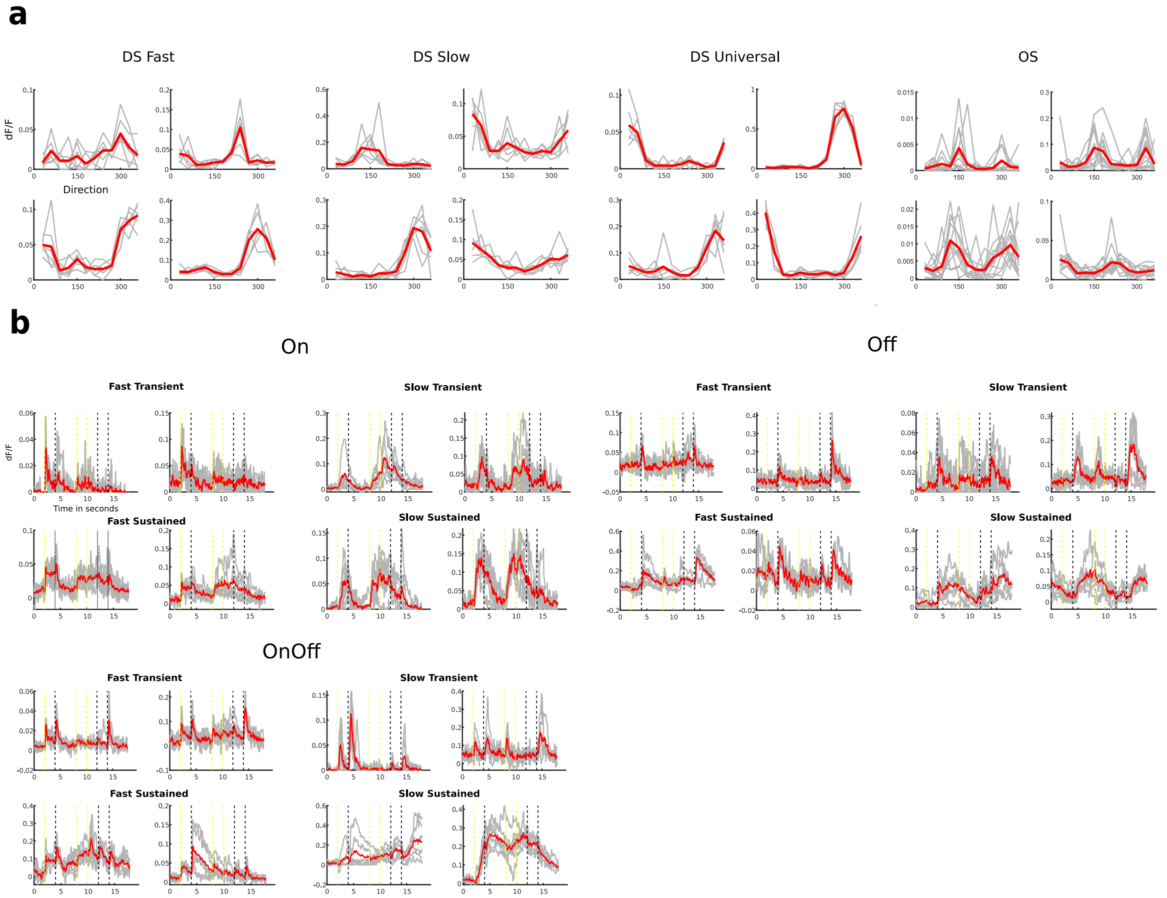


**a)** Response across directions of four randomly chosen cells from each of the OS/DS populations. Grey lines show responses from individual trials while the red lines show the mean response across trials. **b)** Response across time during the ON/OFF phase of the chirp stimulus of two randomly chosen cells from each of the On/Off populations. Grey lines show responses from individual trials while the red lines show the mean response across trials. Dashed yellow line show onset of ON phases, dashed black lines show onset of OFF phases.

##### ***Supplementary figure 6 Relationship between OS/DS and ON/OFF response properties***

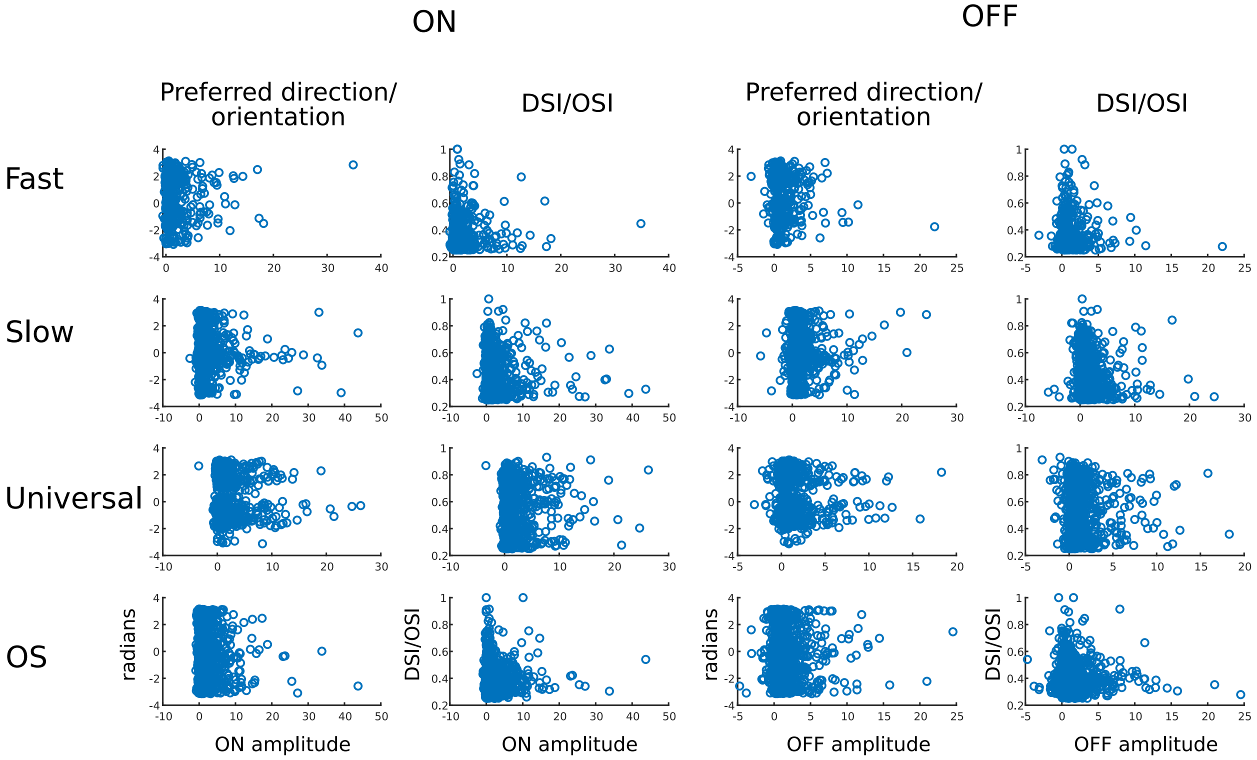


Scatter plots showing relationship between preferred direction/orientation (column 1 and 3) or DSi/OSi (column 2 and 4) and ON (column 1 and 2) or OFF (column 3 and 4) response amplitude (in multiples of baseline standard deviation) across the four OS/DS populations (rows).

#### *Supplementary figure 7 Coincidental overlap of independent data*

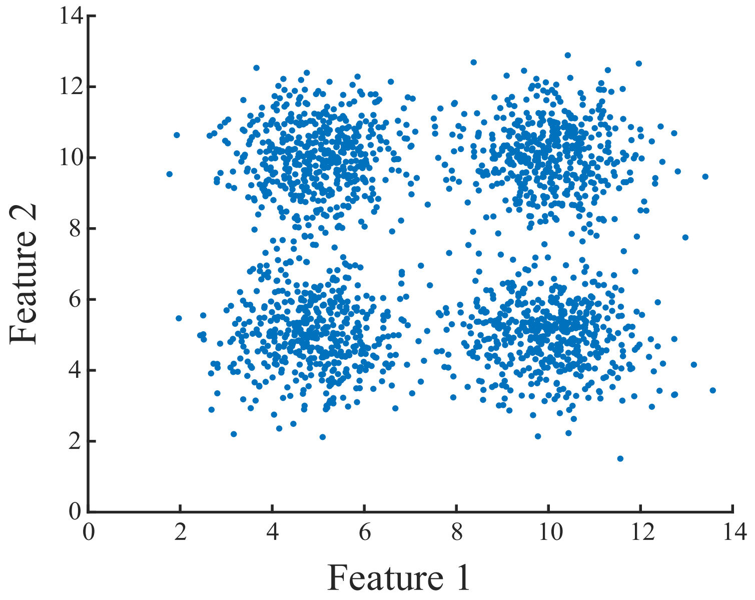


First, we generated two variables, that each consist of two normal distributions with different means and forced them to be independent. We know that variable A holds no information about variable B, and that all overlap between them is coincidental. Nonetheless, if we plot a 1000 simulated cells with variable A along the x-axis, and variable B along the y-axis, a clustering algorithm will identify four clearly separated groups. Why this is problematic can be exemplified using paint with two qualities; variable A is the dry time that can be either fast or slow, variable B is the color. The dry time and the color are completely independent and do not affect one another. It is meaningful to categorize the paint on both qualities, but as the overlap between the two are coincidental all combinations in the matrix made from the two variables will exist. This might be reasonable with two dimensions but with an increasing number of dimensions this becomes increasingly problematic, as the number of clusters will increase by $x^d$, with x being the number of subdivisions within each dimension and d being the number of independent dimensions.

#### *Supplementary figure 8 Overlap between On/Off and OS/DS subtypes*

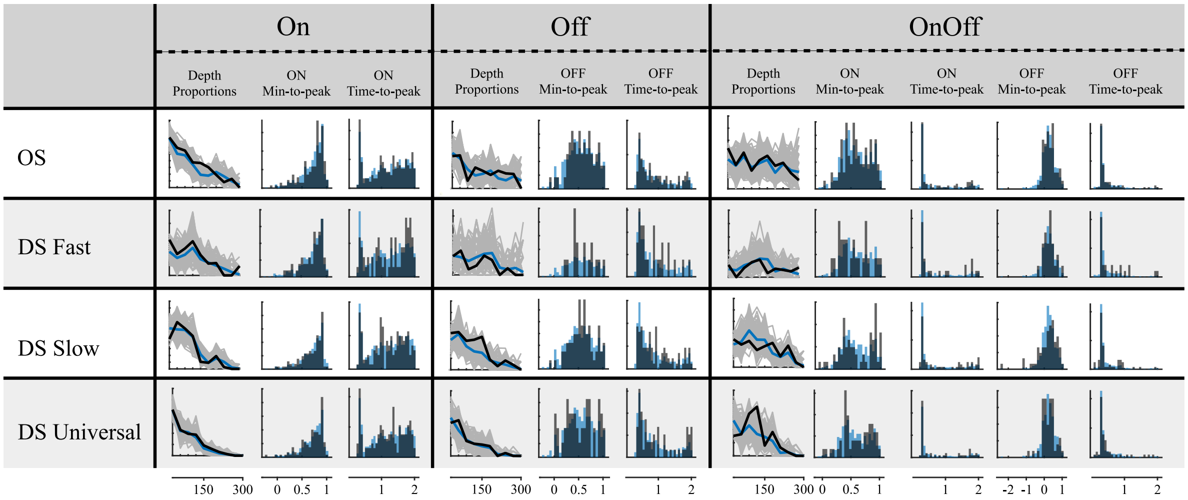


Table showing depth proportions, min-to-peak ratio, and time-to-peak for all subtype combinations. Depth proportions shown as in Figure 1e. Min-to-peak ratio and time-to-peak shown as normalized histograms comparing the distribution of the combined group in dark gray (e.g. On-OS) vs the remaining cells in the parent group in blue (e.g. On-nonOS). x-axis left to right: depth in µm, min-to-peak ratio, and time-to-peak in seconds. y-axis proportion of cells. For OnOff cells min-to-peak ratio and time-to-peak are measured during both the ON and OFF phases.

#### *Supplementary figure 9A Clustering Procedure - Chirp*

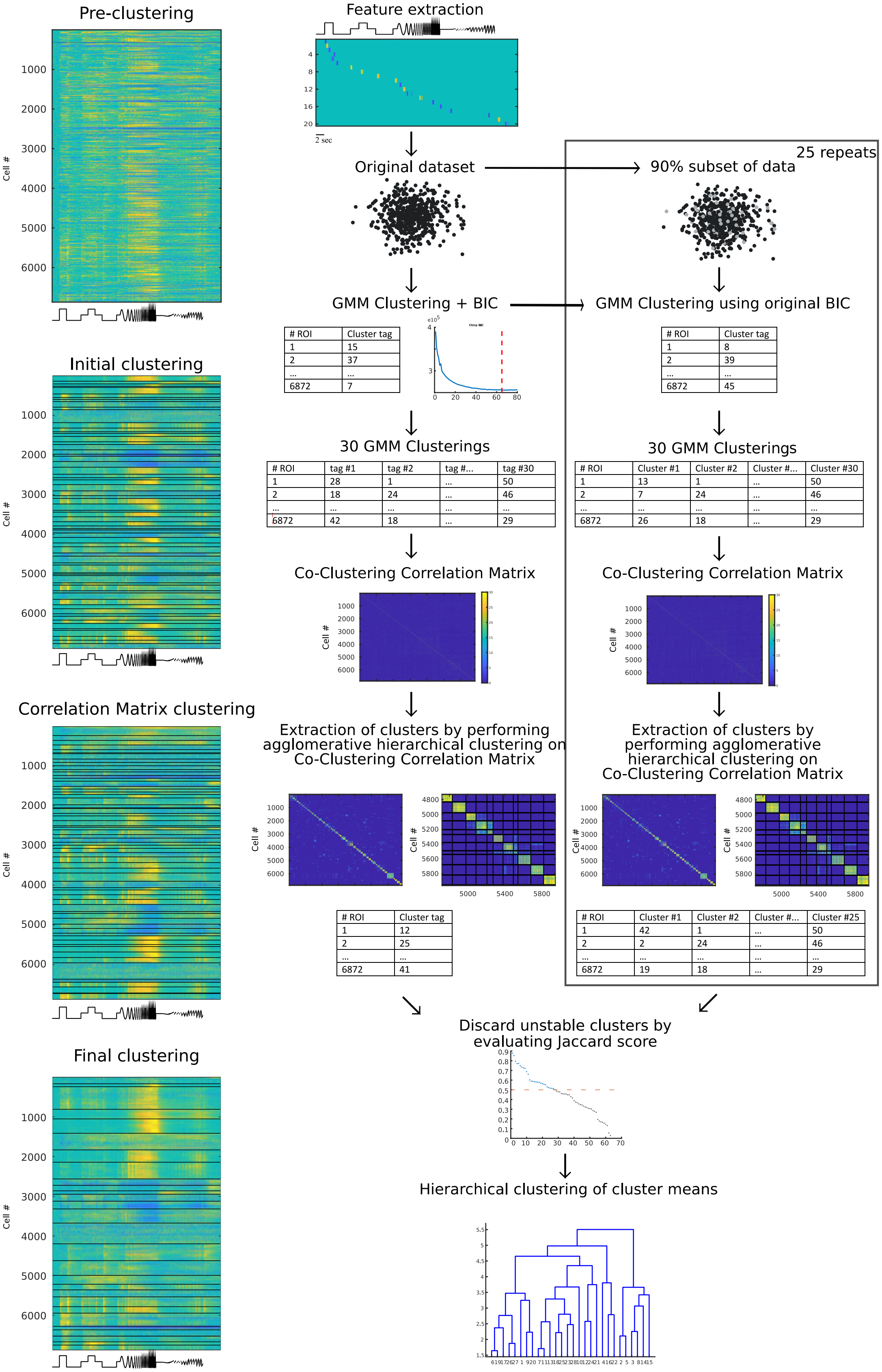


Clustering was performed using Gaussian Mixture Modeling (GMM). The number of clusters were chosen based on calculations of the Bayesian Information Criteria (BIC) on ten preliminary GMM clusterings. Based on an additional 30 GMM clusterings using the number of clusters indicated by the BIC, we constructed a Co-Clustering Correlation Matrix. This matrix was then sorted by performing agglomerative hierarchical clustering, defining new clusters based on cells' co-clustering score. We then used the same clustering procedure on 25 90% subsets and finally calculated the Jaccard Similarity between each cluster from the original data and the most similar cluster in each of the 25 subsample clusters. Clusters with a mean Jaccard similarity score of less than 0.5 were discarded and cells from discarded clusters were re-assigned to the cluster with which it had the highest average co-clustering score. (Schwartz and Yonehara n.d.)

#### *Supplementary figure 9B Clustering Procedure - Drift*

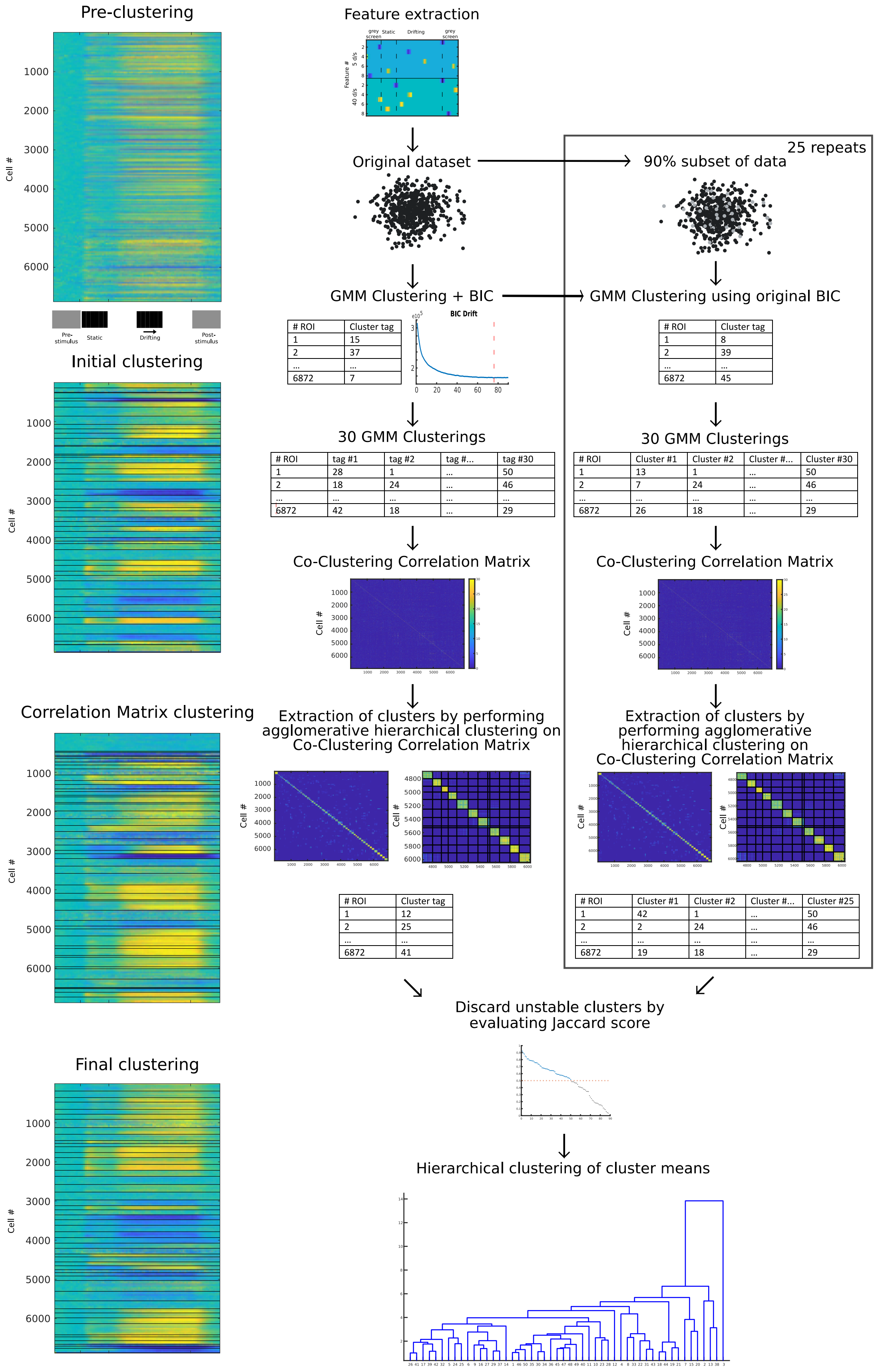


Clustering was performed using Gaussian Mixture Modeling (GMM). The number of clusters were chosen based on calculations of the Bayesian Information Criteria (BIC) on ten preliminary GMM clusterings. Based on an additional 30 GMM clusterings using the number of clusters indicated by the BIC, we constructed a Co-Clustering Correlation Matrix. This matrix was then sorted by performing agglomerative hierarchical clustering, defining new clusters based on cells' co-clustering score. We then used the same clustering procedure on 25 90% subsets and finally calculated the Jaccard Similarity between each cluster from the original data and the most similar cluster in each of the 25 subsample clusters. Clusters with a mean Jaccard similarity score of less than 0.5 were discarded and cells from discarded clusters were re-assigned to the cluster with which it had the highest average co-clustering score. (Schwartz and Yonehara n.d.)

### *Supplementary figure 9C Clustering Procedure - Drift-Chirp*

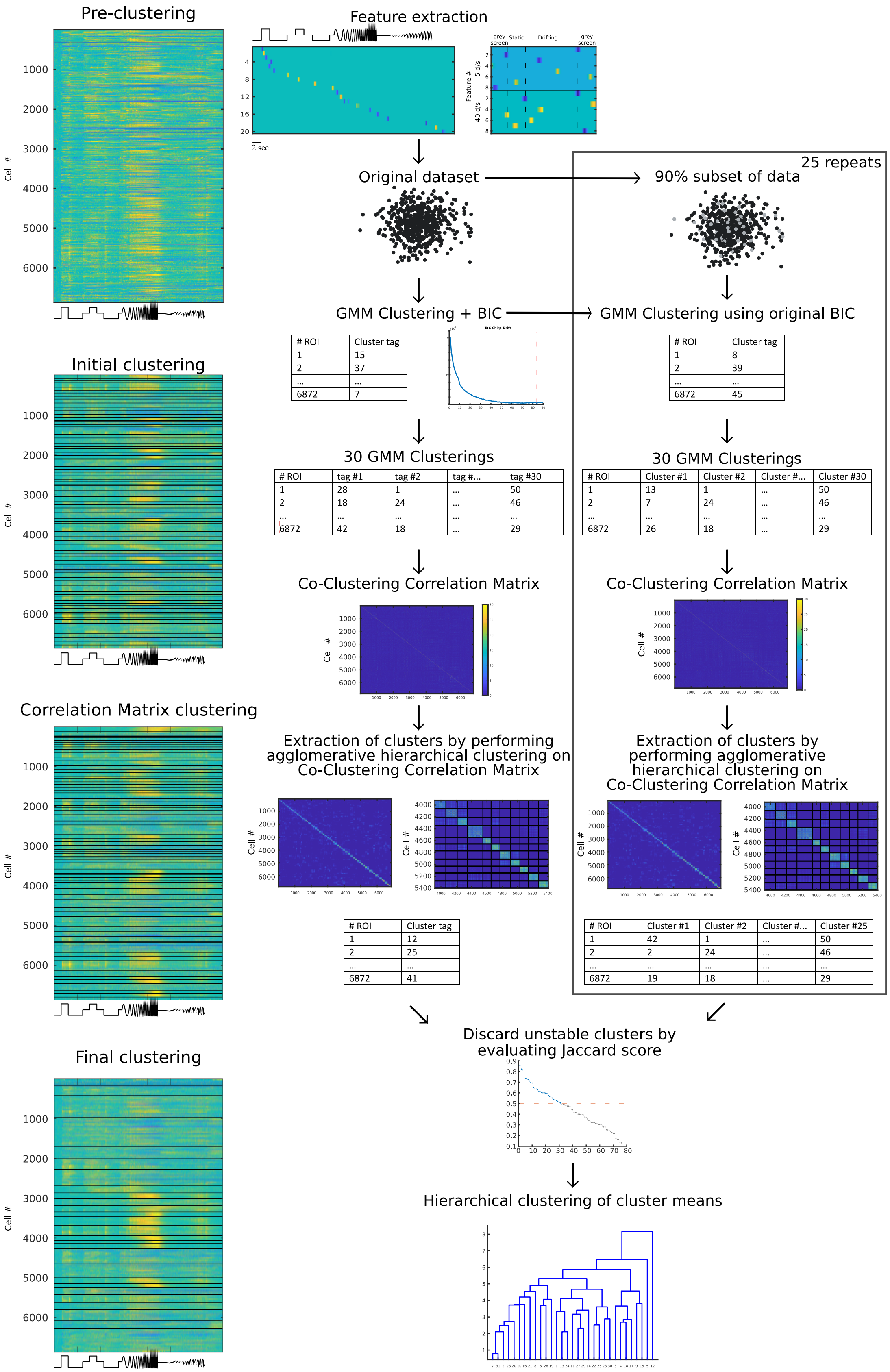


Clustering was performed using Gaussian Mixture Modeling (GMM). The number of clusters were chosen based on calculations of the Bayesian Information Criteria (BIC) on ten preliminary GMM clusterings. Based on an additional 30 GMM clusterings using the number of clusters indicated by the BIC, we constructed a Co-Clustering Correlation Matrix. This matrix was then sorted by performing agglomerative hierarchical clustering, defining new clusters based on cells' co-clustering score. We then used the same clustering procedure on 25 90% subsets and finally calculated the Jaccard Similarity between each cluster from the original data and the most similar cluster in each of the 25 subsample clusters. Clusters with a mean Jaccard similarity score of less than 0.5 were discarded and cells from discarded clusters were re-assigned to the cluster with which it had the highest average co-clustering score. (Schwartz and Yonehara n.d.)

#### *Supplementary figure 10A Cluster Responses - Chirp*

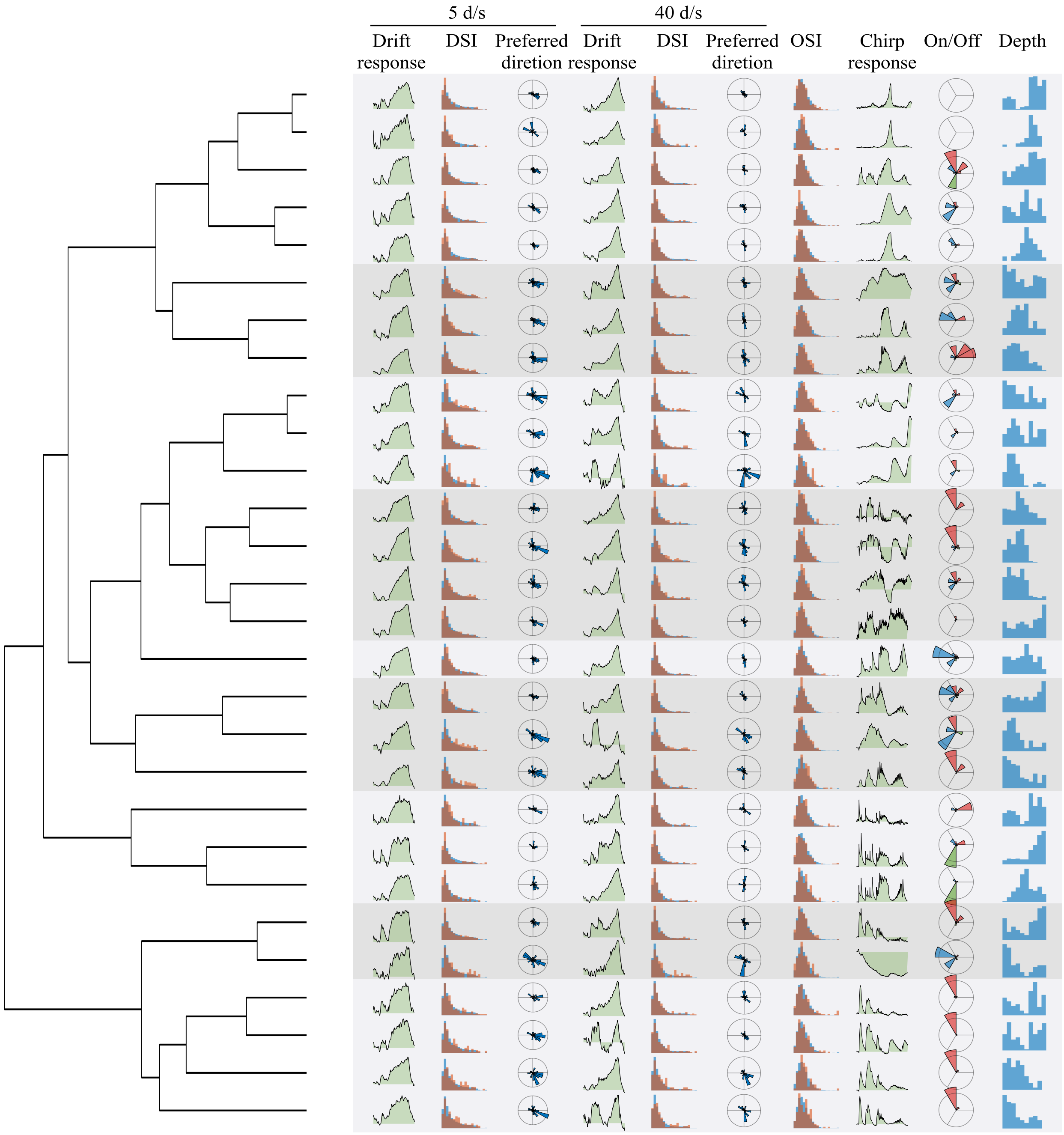


Left: dendrogram for the chirp clustering. Right: mean responses and descriptive statistics for each of the chirp clusters. From left to right the shown parameters are: 1. Drift response: Mean amplitude of the first component of the SVD of the response to the slow grating, 2. DSI: Histogram of the DSi of the response to the slow grating, 3. Preferred direction: Polar histogram of the preferred direction to the slow grating scaled by the proportion of cells in the cluster that were DS to the slow grating, 4-6. Same as 1-3 but for the fast grating, 7. OSi: Histogram of the OSi, 8. Chirp response: Mean response to the chirp stimulus, 9. On/Off: On/Off subtype composition visualized as a polar histogram with red indicating On, blue indicating Off, and green indicating OnOff responses. Within each color there is a further clockwise subdivision into Fast-Transient, Slow-Transient, Fast-Sustained, and Slow-Sustained subtypes, 10. Depth: Histogram showing depth distribution. (Schwartz and Yonehara n.d.)

#### ***Supplementary figure 10B Cluster Responses - Drift***

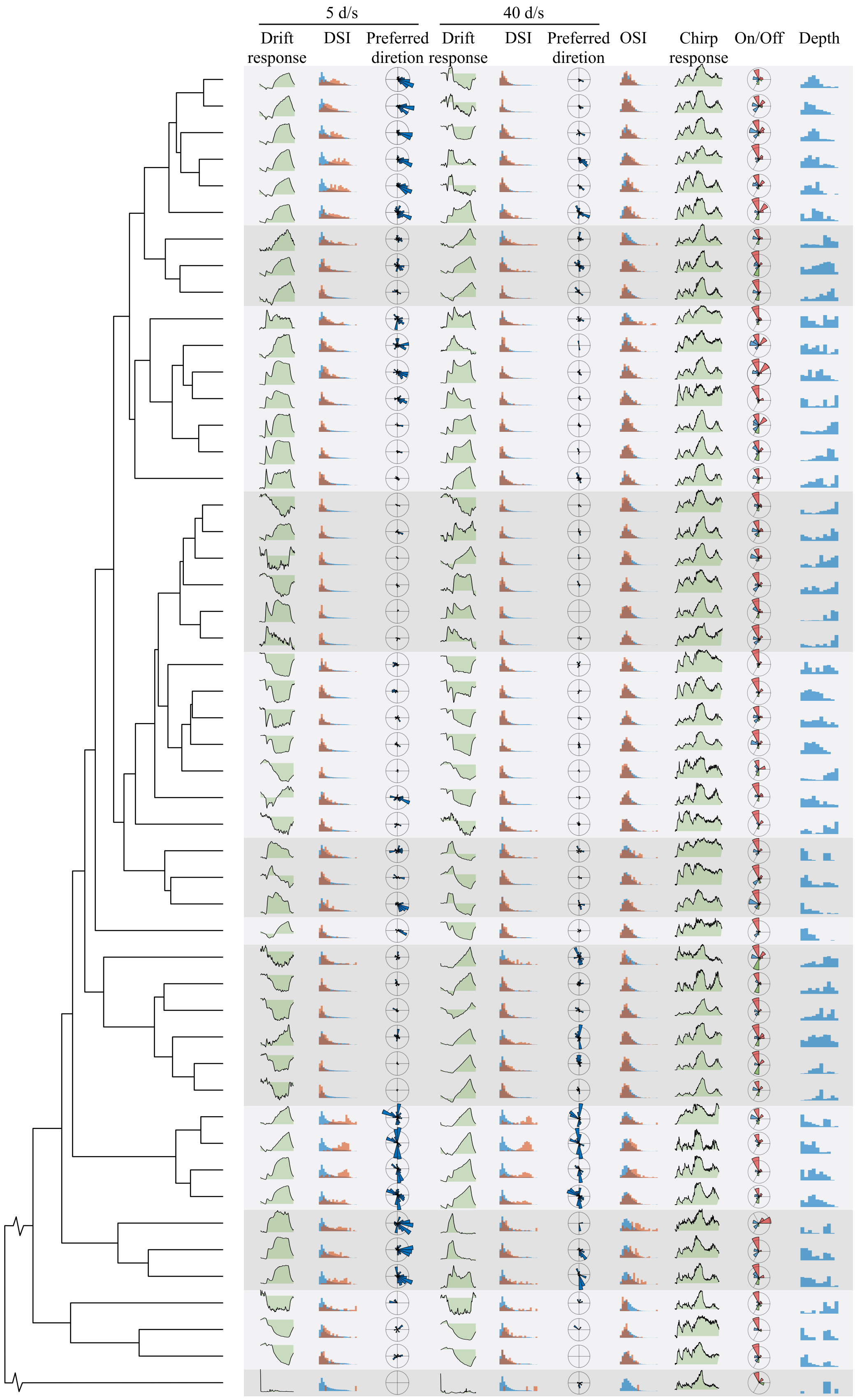


This figure is the same as figure 15A but shows responses and statistics for the Drift clustering. [(Schwartz and Yonehara n.d.)](#)

#### *Supplementary figure 10C Cluster Responses – Chirp-Drift*

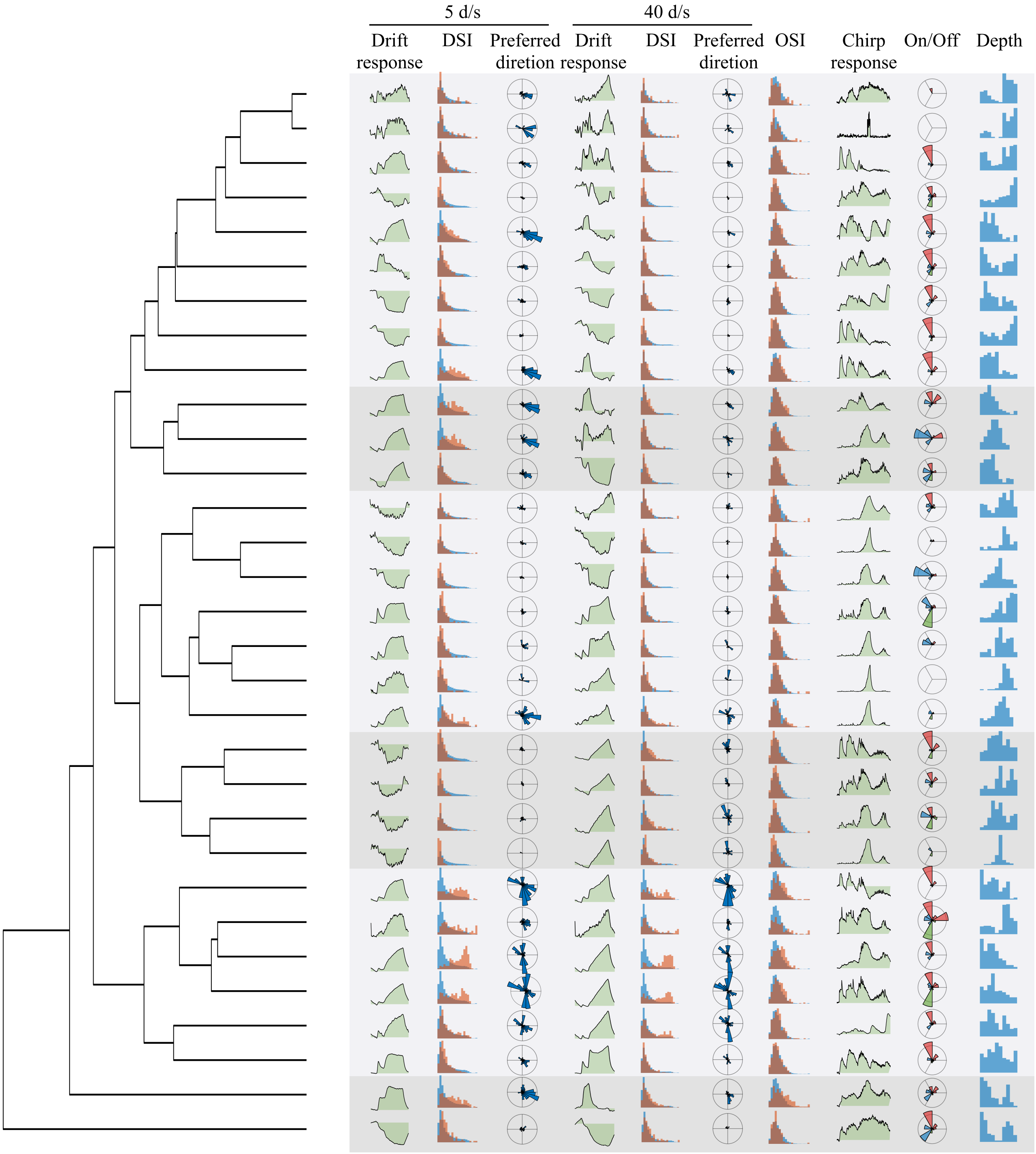


This figure is the same as figure 15A but shows responses and statistics for the Chirp-Drift clustering. [(Schwartz and Yonehara n.d.)](#)

#### ***Supplementary figure 11 Clustering Response Type Separation***

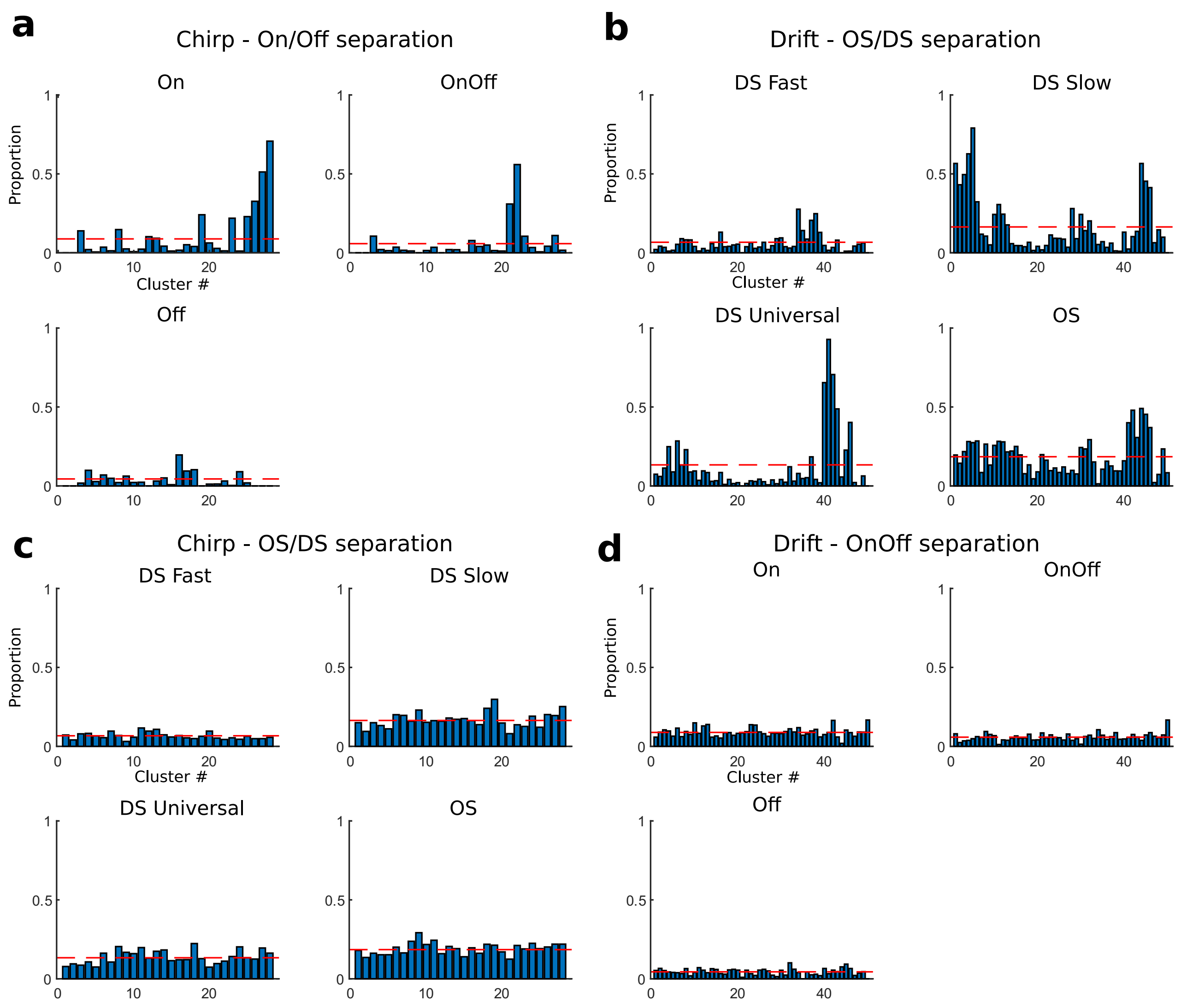


**a)** Proportions of On (top left), OnOff (top right), and Off (bottom) cells across Chirp clusters with dashed red line indicating expected proportion if distribution is random. **b)** Proportions of DS Fast (top left), DS Slow (top right), DS Universal (bottom left), and OS (bottom right) cells across Drift clusters. **c)** Proportions of DS Fast (top left), DS Slow (top right), DS Universal (bottom left), and OS (bottom right) cells across Chirp clusters. **d)** Proportions of On (top left), OnOff (top right), and Off (bottom) cells across Drift clusters.

#### *Supplementary figure 12 Effect of added noise vs added orthogonal features*

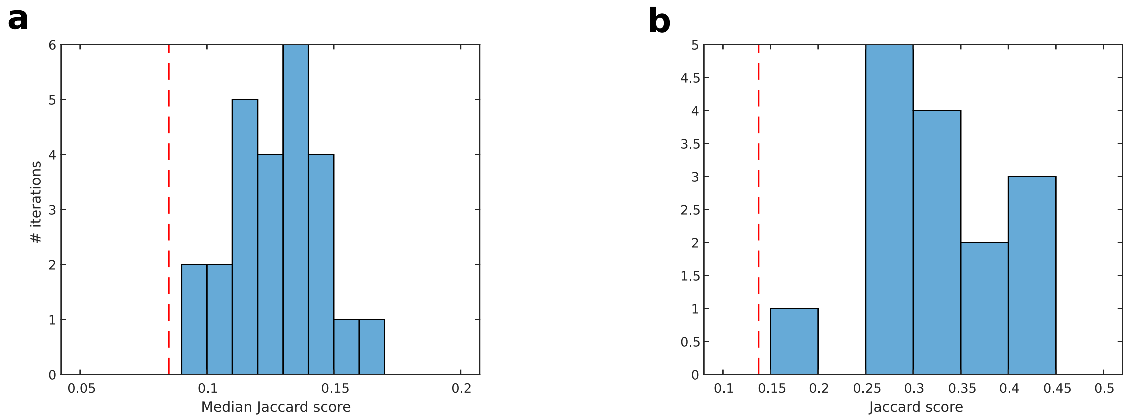


**a)** Median Jaccard score of the chirp clustering vs the chirp-drift clustering (red dashed line) shown on top of a histogram of the median Jaccard scores of the chirp clustering vs chirp-noise clusterings (see methods).
**b)** same as a) using the drift clustering as outset.

***Supplementary figure 13 Population speed spread***

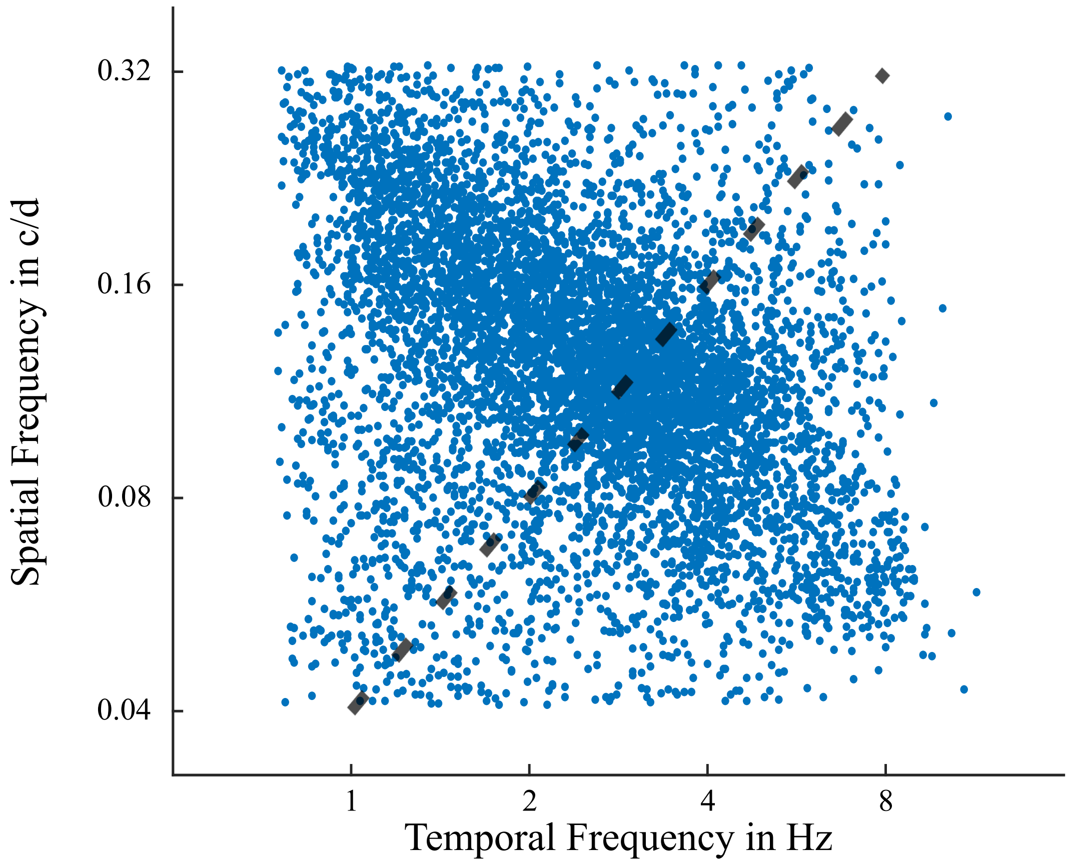


A scatter plot of the preferred spatial frequency of SC cells (blue dots) as a function of their preferred temporal frequency shows a tendency for cells to be placed along on orthogonal to the 25 degrees pr second speed isoline (dashed black line).

*Supplementary figure 14 Identification of SGS-SO border*

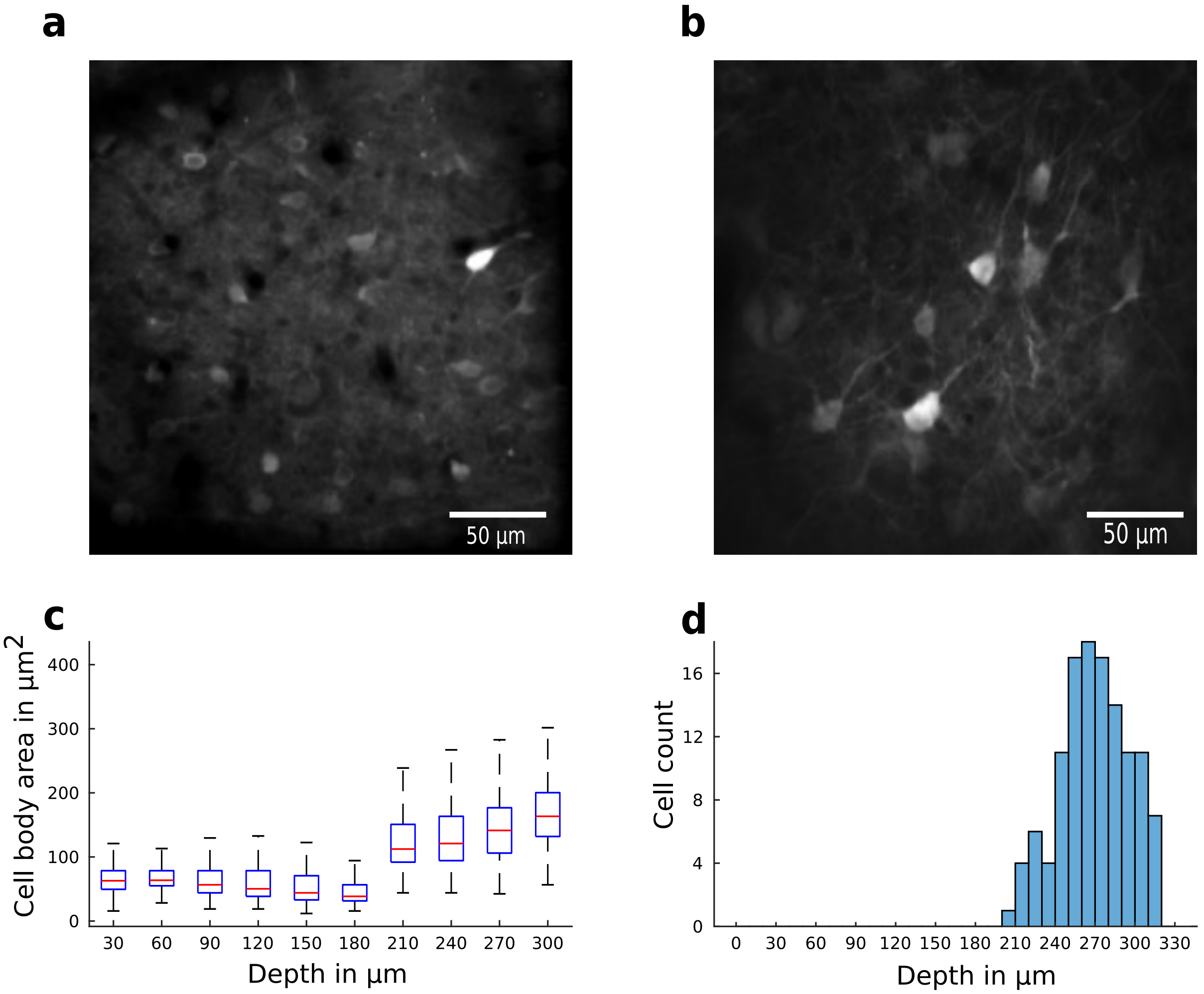


**a)** Image of cells taken 60 µm below SC surface. **b)** Image of cells taken 240 µm below SC surface. **c)** Boxplot showing how cell body size changes across depth. **d)** Histogram showing depth distribution of ntsr1 cells.

#### *Supplementary figure 15 ON OFF Population Responses vs Cutoff*

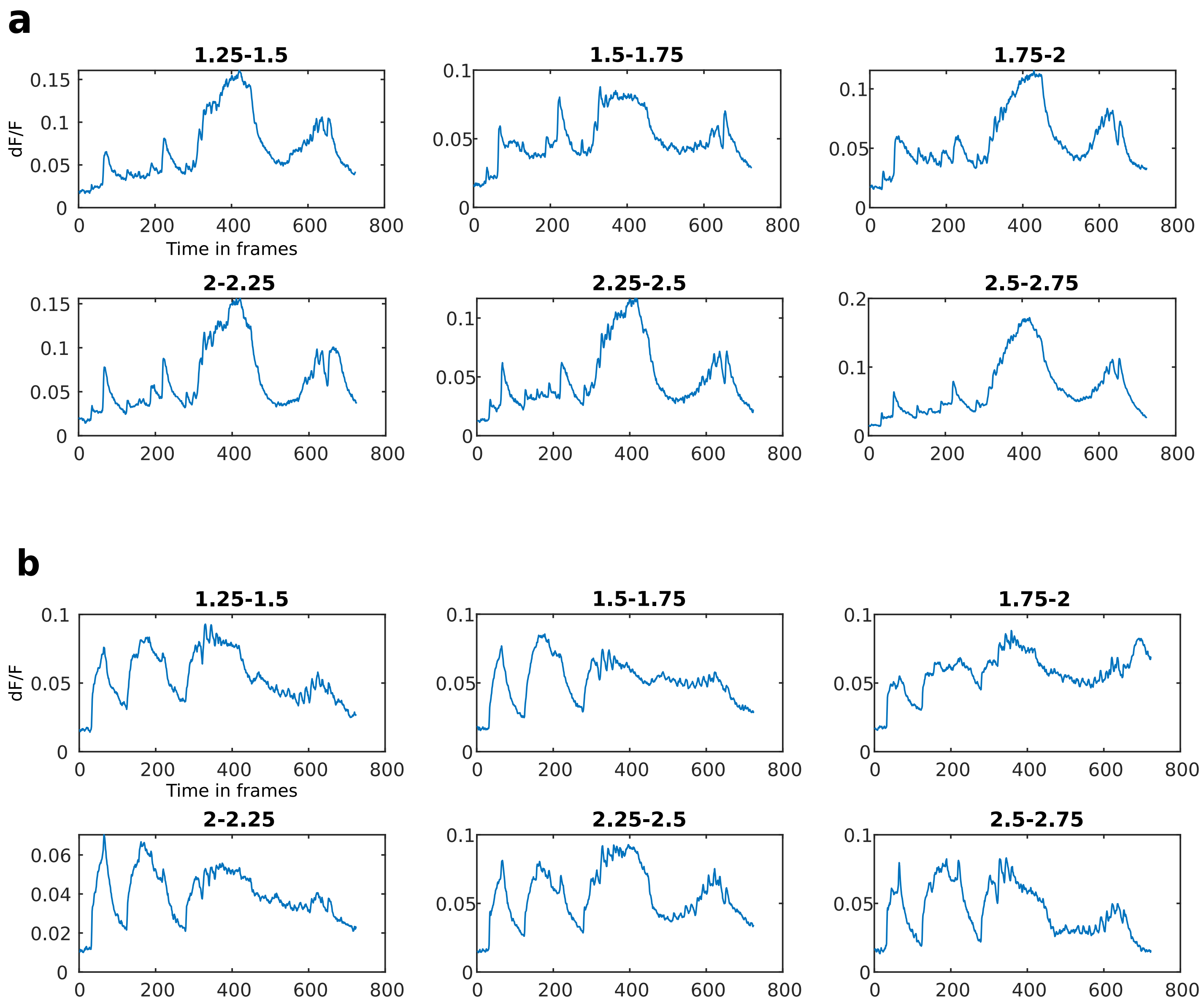


**a)** Plots showing the mean response of cells with an OFF response more than 2.75 times higher than baseline standard deviation and an ON response amplitude within the limits shown above the figure. **b)** Plots showing the mean response of cells with an ON response more than 2.75 times higher than baseline standard deviation and an OFF response amplitude within the limits shown above the figure.

### *Supplementary figure 16 On/Off Subtype Definitions*

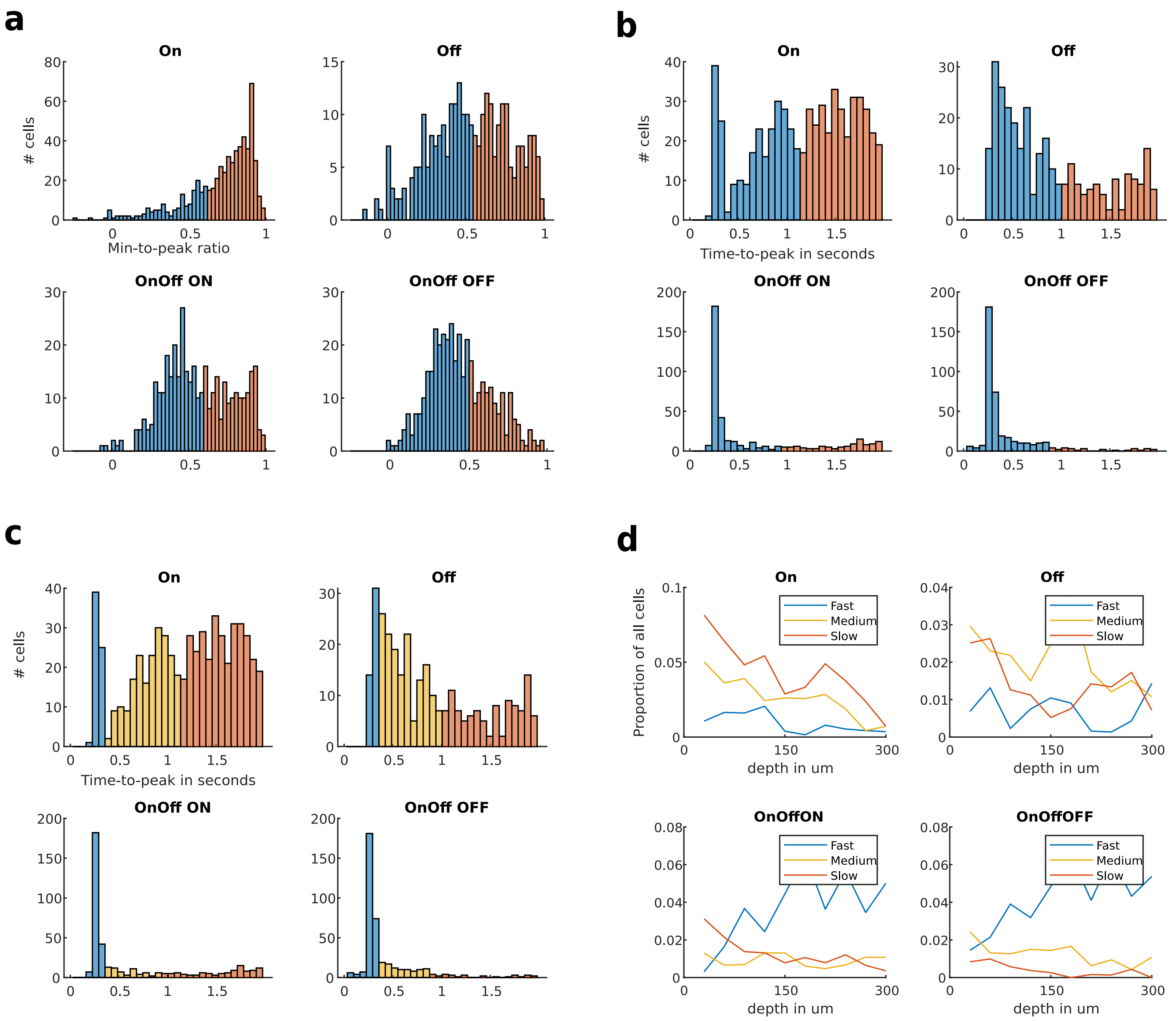


**a)** Histograms of min-to-peak ratio showing k-means based separation into Sustained (red) and Transient (blue) subtypes. **b)** Histograms of time-to-peak showing k-means based separation into Slow (red) and Fast (blue) subtypes. **c)** Same as b) with a further subtype (Medium, yellow) defined based on visual inspection. **d)** Plots of the depth proportions of Fast, Medium, and Slow subtypes for On, Off, OnOff-ON, and OnOff-OFF responses. (Schwartz and Yonehara n.d.)

#### *Supplementary figure 17 Clustering method applied to randomized data*

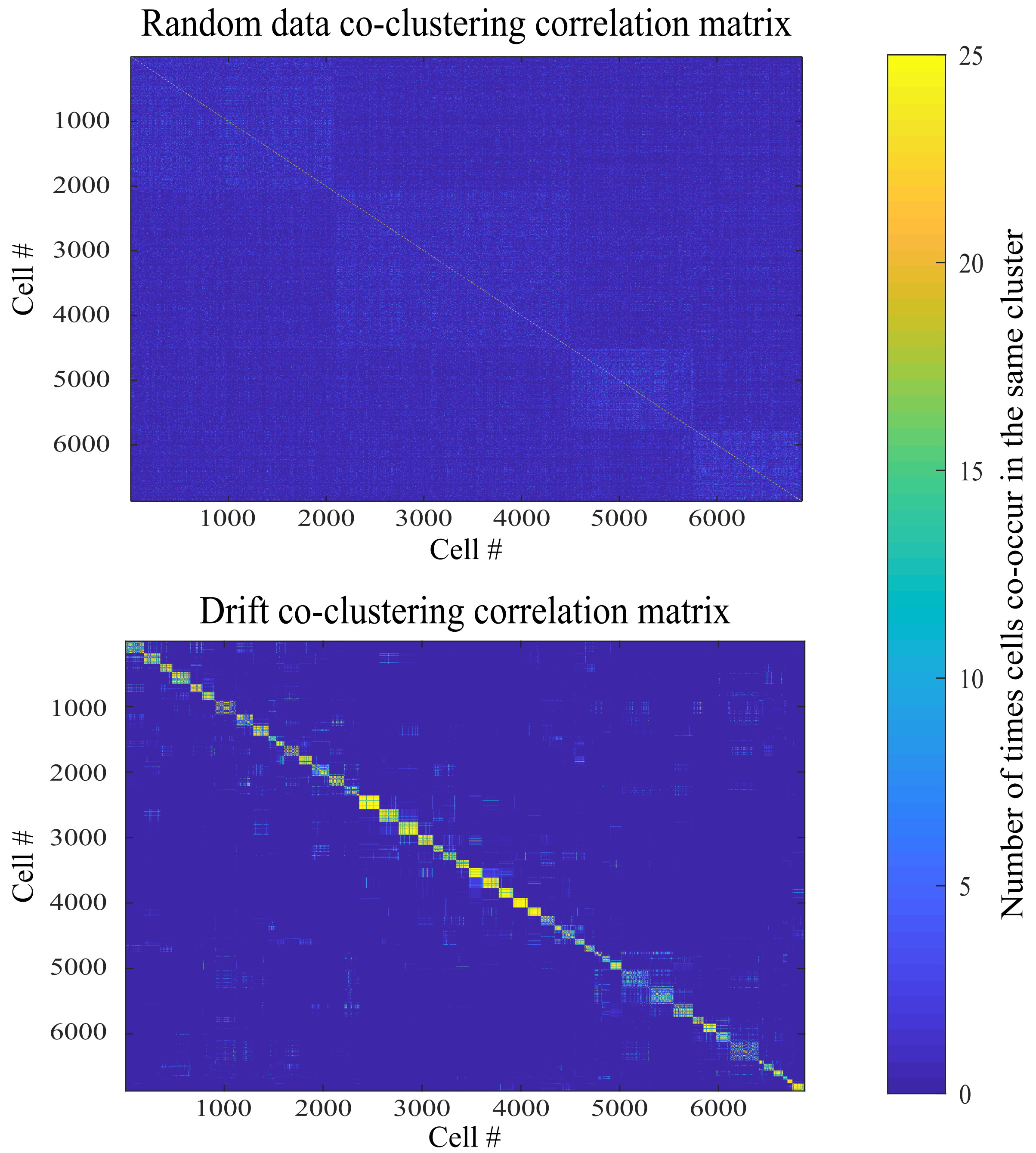


A co-clustering correlation matrix shows how often a pair of cells end up in the same cluster across repeats. The matrix based on random data shows four clusters with low scores (top), whereas the matrix based on actual data shows many highly coherent clusters (bottom).

#### *Supplementary figure 18A Batch Effects - Chirp*

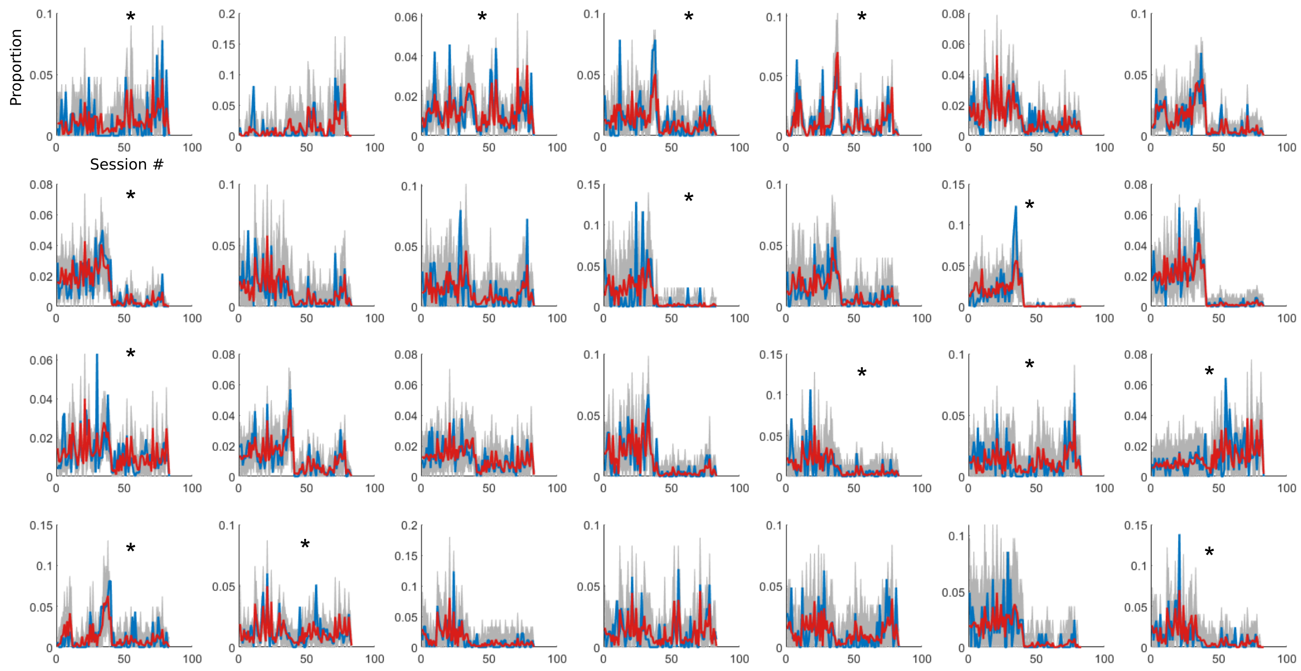


Plots showing proportion of cells stemming from a specific session for each of the 28 clusters from the chirp clustering. Actual proportion shown in red, expected proportion shown in blue, and proportions of 500 random permutations shown in grey. * marks clusters where the actual proportion deviates more than chance from the expected proportion. In some clusters a clear shift in proportion is visible around session 40. The cause of this shift is that all sessions before 40 were recorded from the stratum griseum superficiale (SGS) while those after session 40 were recorded from the stratum opticum (SO). (Schwartz and Yonehara n.d.)

#### *Supplementary figure 18B Batch Effects - Drift*

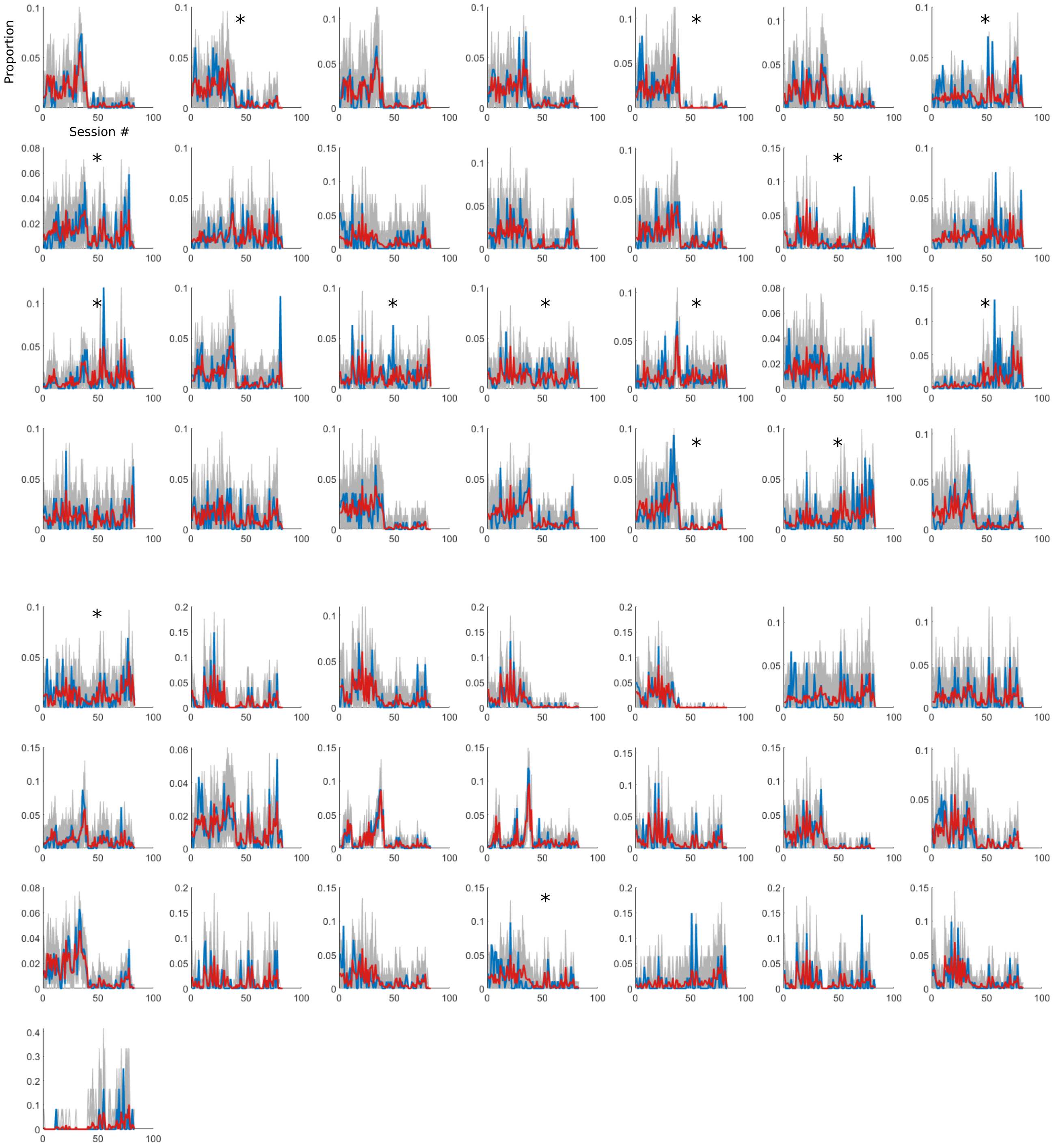


Plots showing proportion of cells stemming from a specific session for each of the 50 clusters from the drift clustering. Actual proportion shown in red, expected proportion shown in blue, and proportions of 500 random permutations shown in grey. * marks clusters where the actual proportion deviates more than chance from the expected proportion. In some clusters a clear shift in proportion is visible around session 40. The cause of this shift is that all sessions before 40 were recorded from the stratum griseum superficiale (SGS) while those after session 40 were recorded from the stratum opticum (SO). (Schwartz and Yonehara n.d.)
